# Rethinking simultaneous suppression in visual cortex via compressive spatiotemporal population receptive fields

**DOI:** 10.1101/2023.06.24.546388

**Authors:** Eline R Kupers, Insub Kim, Kalanit Grill-Spector

## Abstract

When multiple visual stimuli are presented simultaneously in the receptive field, the neural response is suppressed compared to presenting the same stimuli sequentially. The prevailing hypothesis suggests that this suppression is due to competition among multiple stimuli for limited resources within receptive fields, governed by task demands. However, it is unknown how stimulus-driven computations may give rise to simultaneous suppression. Using fMRI, we find simultaneous suppression in single voxels, which varies with both stimulus size and timing, and progressively increases up the visual hierarchy. Using population receptive field (pRF) models, we find that compressive spatiotemporal summation rather than compressive spatial summation predicts simultaneous suppression, and that increased simultaneous suppression is linked to larger pRF sizes and stronger compressive nonlinearities. These results necessitate a rethinking of simultaneous suppression as the outcome of stimulus-driven compressive spatiotemporal computations within pRFs, and open new opportunities to study visual processing capacity across space and time.

## 2 Introduction

The human visual system has limited processing capacity. We are worse at processing multiple stimuli presented at once than when the identical stimuli are shown one after the other in the same location. This drop in performance has been observed in a variety of visual tasks, such as searching for a target among distractors^1,2^, recognizing an object when surrounded by flankers^3^, or keeping multiple items in short-term visual working memory^4^.

A neural phenomenon attributed to limited visual capacity is simultaneous suppression: A reduced response when multiple visual stimuli are presented at once than when the identical stimuli are shown one after the other in sequence^5–12^. Simultaneous suppression is prevalent and robust: it is observed across visual cortex, from the level of single-neuron spiking^5–7^, all the way to the level of entire visual areas using fMRI^8–12^, with large effect sizes: up to 2-fold amplitude differences between sequential and simultaneous presentations of otherwise identical stimuli^8,9,12^.

A prevailing explanation linking simultaneous suppression to visual capacity is based on the influential theory of biased competition^7,8,13^. This theory argues that visual processing capacity is determined by the computational resources afforded by receptive fields, where the visual system prioritizes inputs that are behaviorally relevant for further processing. When a visual stimulus is presented alone in the receptive field, the item can be fully processed with the limited neural resources. However, when multiple stimuli are presented at once in the receptive field, the stimuli compete for neural resources, resulting in a reduced neural response. This explanation of stimuli competing for neural resources is linked to visual attention, as it has been suggested that competition can be governed by task or behavioral demands^13^. As such, a large body of research has examined how simultaneous suppression is modulated by visual attention^7,8,14–16^ and stimulus context^10,11^. However, it is unknown how stimulus-driven computations within receptive fields may give rise to simultaneous suppression in the first place. Thus, the goal of the present study is to operationalize and elucidate the computational mechanisms underlying simultaneous suppression in human visual cortex.

A key prediction stemming from the biased competition theory is that simultaneous suppression will only occur in neurons whose receptive fields are large enough to encompass several stimuli^9,12^. It is well documented that the size of receptive fields^17^ and population receptive fields (pRFs, aggregate receptive field of the neuronal population in an fMRI voxel^18,19^) progressively increase from earlier to higher areas up the visual hierarchy. Consistent with this idea, several studies reported that simultaneous suppression systematically increases up the visual hierarchy and is absent in V1^7–9,12^, suggesting that the lack of suppression in V1 is because its receptive fields are too small to encompass multiple visual stimuli. However, there is an assumption in prior work, which is that neurons sum inputs linearly over the duration of the stimulus^7,9,12,20^. This assumption has emphasized research on how the spatial overlap between stimuli and the receptive field may affect simultaneous suppression^8,9,12^ and less on how stimulus timing may affect simultaneous suppression.

As stimuli have identical duration, size, and location across simultaneous and sequential conditions, neurons summing linearly over visual space and linearly over stimulus duration are predicted to respond identically in these conditions. Therefore, the lower responses for simultaneous than sequential presentations suggest nonlinear, and in particular, subadditive summation. Although linear summation within receptive fields is observed in some cases^21–23^, violations of response linearity in the human visual system have been extensively reported. Spatially, responses to bigger stimuli are typically less than the sum of responses to smaller stimuli^22,24–27^. Temporally, responses to longer stimuli are typically smaller than the sum of responses to shorter stimuli^28–44^. While both hemodynamic^45–47^ and neural^27,35,39–42,48–51^ mechanisms may contribute to observations made with fMRI, empirical and computational modeling studies suggest that the observed subadditivity is driven by compressive summation of visual inputs within neurons’ receptive fields across space^27,50^ and across time^33,36,38,40–44^. Therefore, rather than considering how context or task demands affect simultaneous suppression, we asked: to what extent is simultaneous suppression a consequence of compressive computations within receptive fields in visual cortex? Here, we consider two possible classes of compressive neural mechanisms that may generate simultaneous suppression: compressive spatial summation and compressive spatiotemporal summation.

Compressive spatial summation within the receptive field predicts that the response to multiple stimuli presented together within the pRF (as in simultaneous conditions) will be lower than the sum of responses to the individual stimuli shown alone (as in sequential conditions). As the duration of stimuli are matched between the simultaneous and sequential conditions, the spatial summation hypothesis predicts that the level of simultaneous suppression will depend only on the spatial overlap between the stimuli and the pRF. As both average receptive field size and compressive nonlinearities increase up the visual hierarchy^27,50^, compressive spatial summation also predicts that the level of simultaneous suppression will increase from earlier to later visual areas.

Compressive spatiotemporal summation within the receptive field predicts that simultaneous suppression will not only depend on the spatial overlap between stimuli and the receptive field, but also the timing of stimuli. This prediction is based on the empirical observation that neuronal responses to visual stimuli vary over time, typically showing an initial strong transient response at stimulus onset (lasting for 100-200 ms), followed by a weaker sustained response lasting for the duration of the stimulus^32,42,44,52–55^, and often a transient response at stimulus offset^42,44,54^. These nonlinear temporal dynamics suggest that presenting all stimuli at once in the pRF (as in simultaneous conditions) results in two transients (at stimulus onset and offset). This response will be lower than the accumulated response induced by multiple visual transients when presenting multiple stimuli one-by-one in rapid fashion (as in sequential conditions). Thus, the spatiotemporal hypothesis predicts that the level of simultaneous suppression will depend on both the spatial overlap between the stimuli and the pRF, and the difference in the number of visual transients between simultaneous and sequential conditions. Similar to compressive spatial nonlinearities, compressive temporal nonlinearities also increase up the visual hierarchy^39–42,44,51^, predicting an increase in the level of suppression as pRF size and spatiotemporal compression increase.

Here, we used fMRI and a computational pRF framework to distinguish between these hypotheses. We conducted two fMRI experiments. In the first (SEQ-SIM, **Fig 1A**), we measured responses to sequentially or simultaneously presented stimuli and examined how stimulus size and timing affect the level of simultaneous suppression in each voxel of the visual system (**Fig 1B**). In the second experiment (retinotopy, **Fig 1D**), we estimated each voxel’s spatial pRF parameters and used estimated parameters in a pRF modeling framework to predict the BOLD time series for each voxel in the SEQ-SIM experiment. We then implemented several pRF models in our modeling framework to computationally test our hypotheses. To test the compressive spatial summation hypothesis, we used a compressive spatial summation (CSS) pRF model^27^ as it successfully predicts subadditive responses to stimuli of different sizes in pRFs across the visual hierarchy. To test the compressive spatiotemporal summation hypothesis, we used a novel compressive spatiotemporal (CST) summation pRF model^51^, which captures pRF responses to a large range of spatial and temporal stimulus conditions by modeling neural responses in units of visual degrees and milliseconds.

**Figure 1.**
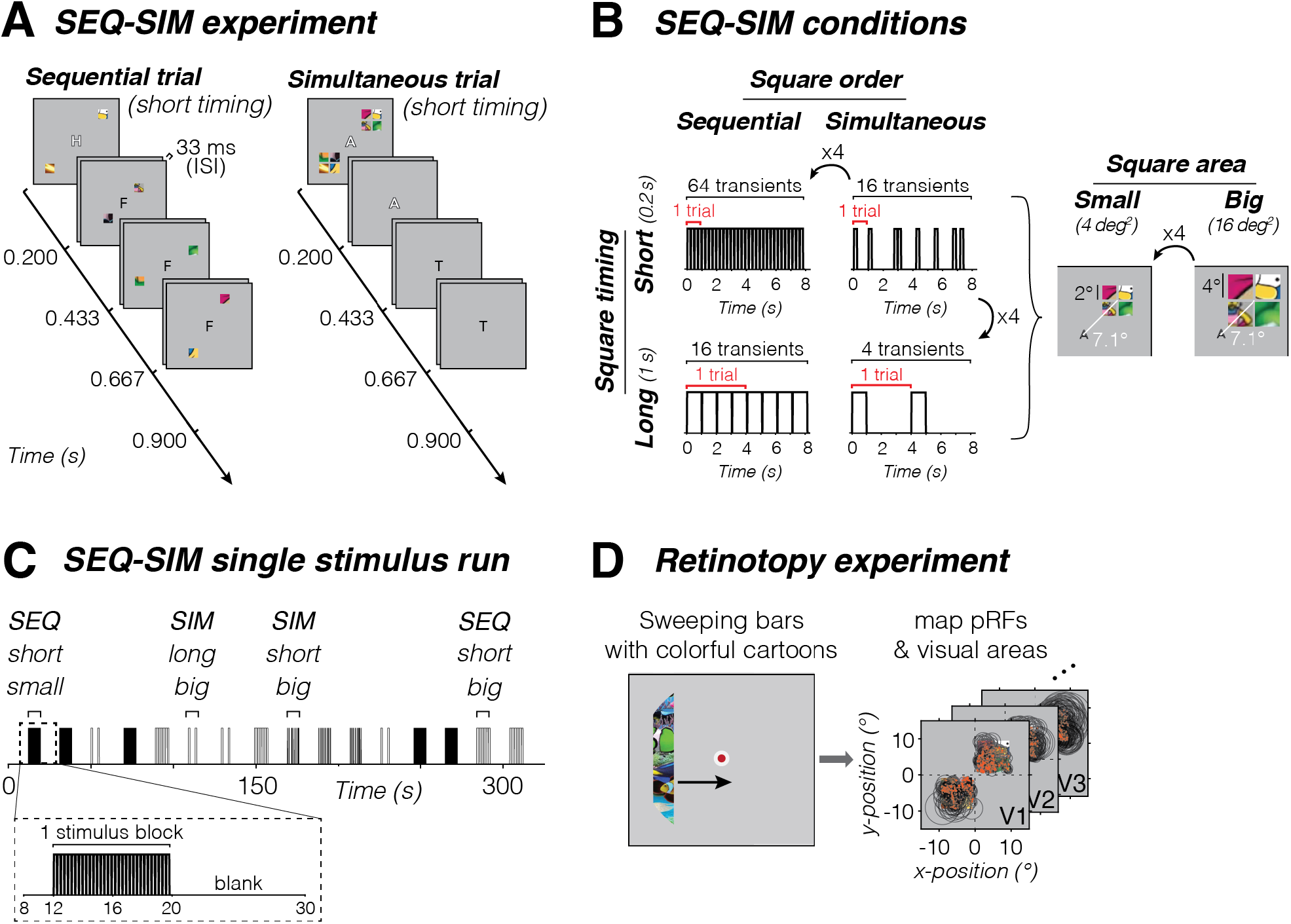
Overview of fMRI experiments. (A) SEQ-SIM experiment. Example trial sequences for small and short stimuli. Four colorful squares were presented in the upper left and lower right quadrants, presented either sequentially in random order (left) or simultaneously (right) interspersed by blank periods to match the trial duration. Observers performed a 1-back RSVP letter task at fixation. Letter is enlarged for visibility. **(B) Stimulus conditions.** Square stimuli were shown in one of two presentation timings (0.2 s or 1 s) and in one of two sizes (4 deg^2^ or 16 deg^2^). Number of trials per block was adjusted to create a 4:1 ratio in number of transients (stimulus onsets or offsets) for short vs long durations. The number of transients indicated is based on a pRF overlapping all four squares, e.g., for 1-s sequentially-presented squares there are 16 transients per block: 4 stimulus frames x 2 on/offsets x 2 trials. If a pRF overlaps only a single square (time course not shown), the number of transients will be identical for SEQ and SIM pairs. For each SEQ-SIM pairing, individual squares were shown for the same duration within a single trial (red bracket). Trials were repeated within an 8-s block (black bracket), where square content was updated for each trial. **(C) Example of SEQ-SIM stimulus run.** A single 332-s SEQ-SIM run contained 16 8-s pseudo-randomized stimulus blocks, interspersed with 12-s blank periods. Data are analyzed in 23-s time windows containing pre-stimulus baseline period, one stimulus block, and subsequent blank period (zoom). **(D) Retinotopy experiment.** Observers viewed bars containing cropped cartoon stimuli traversing the visual field (left, Toonotopy^56^) while fixating and performing a color change detection task at fixation^56^. Data were used to define visual areas and select pRFs with centers overlapping stimulus quadrants in the main experiment (right). Fixation dot is enlarged for visibility.

## 3 Results

To investigate what factors affect simultaneous suppression, we designed an fMRI experiment in which participants viewed colorful patterned square stimuli in upper and lower quadrants while performing a 1-back RSVP letter task at fixation. Squares could either be presented sequentially (one after the other, in random order) or simultaneously (all at once) (**Fig 1A**). For each pair of sequential and simultaneous conditions, individual square presentation is identical in size and duration within an 8-s block such that linear summation of visual inputs in space and time will generate identical responses for both sequence types. To distinguish between spatial and spatiotemporal mechanisms of simultaneous suppression, we varied square size and timing (**Fig 1B,C**). Additionally, participants completed an independent retinotopy experiment^56^ to delineate visual areas and estimate spatial pRF parameters in each voxel (**Fig 1D**).

In each visual area, we measured BOLD responses in voxels which pRF centers overlapped the quadrants with SEQ-SIM stimuli. We then determined how spatial and temporal stimulus properties affect simultaneous suppression for each pRF across visual areas spanning ventral, lateral, and dorsal processing streams. We predict that if simultaneous suppression is of spatial origin, there will be greater suppression in higher-level than early visual areas because those higher-level areas contain larger pRFs that will overlap multiple squares and also show greater spatial compression^27^. Additionally, we predict that varying square size but not timing will affect simultaneous suppression. If simultaneous suppression is of spatiotemporal origin, in addition to observing greater suppression for larger pRFs in higher-level areas, we also predict stronger suppression for long (1 s) than short (0.2 s) presentations because the former has longer sustained stimulus periods, resulting in four times fewer visual transients in the 8-s stimulus blocks than the latter (**Fig 1B**).

To give a gist of the data, we first show results from example voxels in early (V1) and higher-level (VO1/2) areas of the ventral stream. These visual areas differ in overall pRF size and spatial compression: V1 pRFs are small and typically overlap only one square, and VO1/2 pRFs are large and overlap multiple squares.

### 3.1 V1 voxels with small pRFs show modest to no simultaneous suppression

For a single V1 voxel with a small pRF overlapping only a single square, we find similar responses for simultaneous vs sequential presentations for the two stimulus sizes and presentation timings (**Fig 2A**) as the stimulus within the pRF is identical across the two types of presentation sequences. In other words, this voxel shows no simultaneous suppression. Additionally, we observe that for this V1 voxel responses are larger for short presentations (many visual transients) vs long presentations (few visual transients) even though the total duration of stimulation across the blocks is identical. However, there is no difference in the response amplitude for small vs big squares of the same duration (left vs right panels).

**Figure 2.**
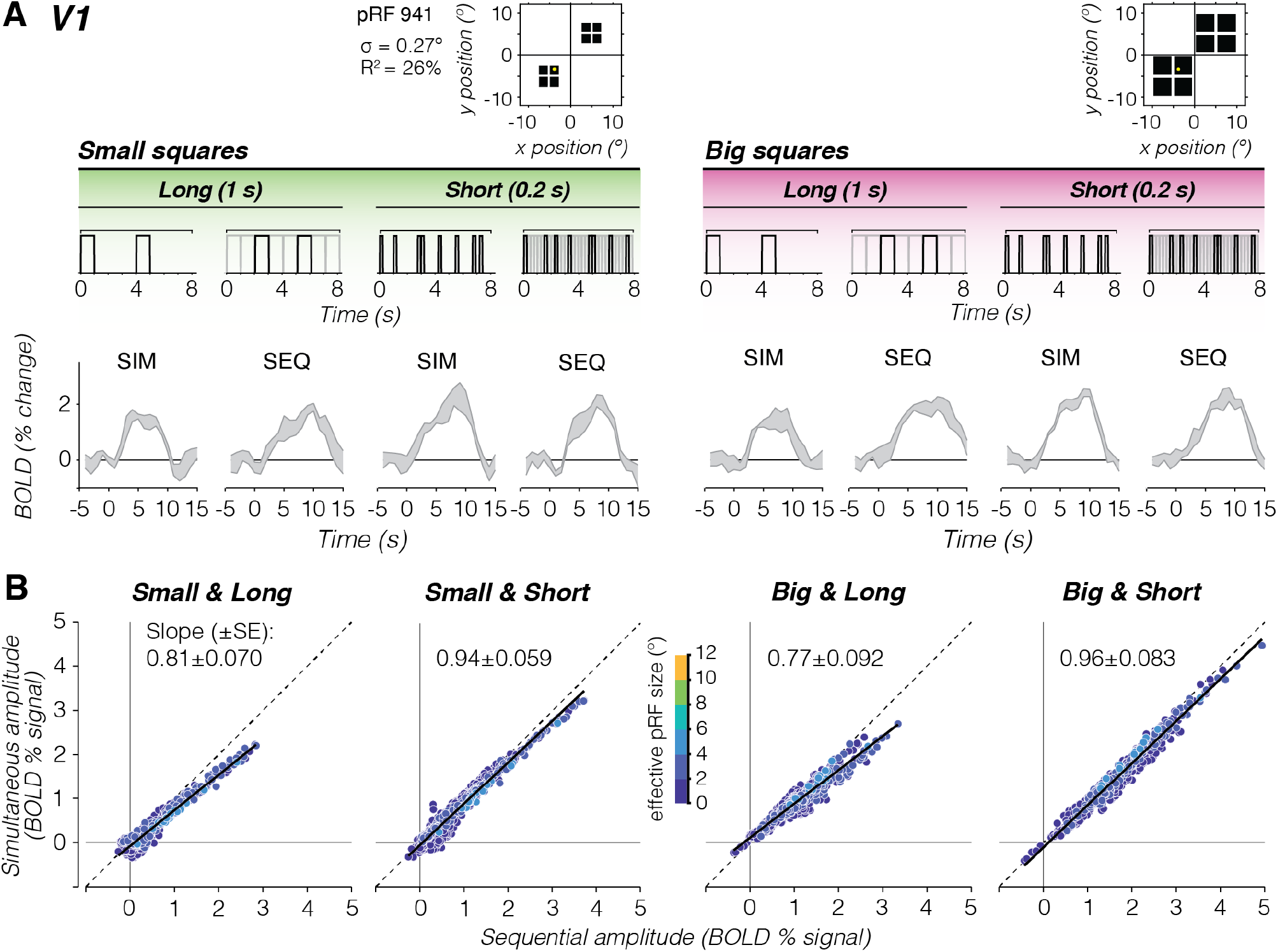
V1 voxels show no to little simultaneous suppression. **(A) Example V1 voxel with small pRF overlapping a single square.** The example voxel’s pRF (yellow circle) is superimposed on square locations (black). Gray time courses show the example voxel’s average BOLD time series ± SEM across block repeats for each stimulus condition. Above each time series is an example stimulus sequence for each condition in an 8-s block. *Gray sequence:* time course including all square stimuli. *Black sequence:* time course for small pRF overlapping one square. **(B) Relation between BOLD amplitude (% signal) for simultaneous vs sequential blocks, for each size/duration condition.** Data include all V1 voxels from participant S3 with pRFs overlapping square stimuli, averaged across a 9-s time window centered on the peak response. Each dot is a voxel, colored by effective pRF size from the retinotopy model fit (σ/√CSS_n_). *Dashed line:* No suppression. *Solid black line:* Linear mixed model (LMM) line fit for this participant’s V1 data. Slope (±SE) across voxels are indicated in each panel.

To assess simultaneous suppression, we compare single voxel response amplitudes for simultaneous vs sequential presentations for a given stimulus condition. No suppression will result on voxels falling on the identity line, whereas simultaneous suppression will result in voxels below the diagonal. In V1, we find that many voxels fall closely or just below the identity line (**Fig 2B**, example participant; **Supplementary Fig 1**, all participants) even as response levels were higher for short vs long stimulus presentation timings. To quantify this relationship, we fit a linear mixed model (LMM) relating the simultaneous amplitude to the sequential amplitude across V1 voxels using a fixed interaction effect for conditions, and a random effect for participants (i.e., intercepts and slopes vary per participant and condition, **Equation 10**). LMM slopes of 1 indicate no suppression, slopes less than 1 indicate simultaneous suppression, where smaller slopes correspond to stronger suppression levels.

Across participants, the LMM captures 86% of the variance in V1, with the following average (± SEM) suppression levels: small and long squares: 0.81±0.069 (CI_95%_=0.56–1.06), small and short squares: 0.85±0.058 (CI_95%_=0.73–0.96), big and long squares: 0.85±0.090 (CI_95%_=0.56–1.4), and big and short squares: 0.84±0.081 (CI_95%_=0.57–1.1). Thus, V1 voxels with relatively small pRFs show modest to no simultaneous suppression.

### 3.2 Strong simultaneous suppression for large pRFs in higher-level visual areas

For a single VO voxel with a large pRF overlapping all four large squares, we find lower responses for simultaneous than sequential presentations for both square sizes and presentation timings (**Fig 3A**). In other words, this voxel shows simultaneous suppression across all experimental conditions. Additionally, we observe that the overall response amplitudes of this voxel are larger for the big squares and short presentations compared to the small squares and long presentations.

**Figure 3.**
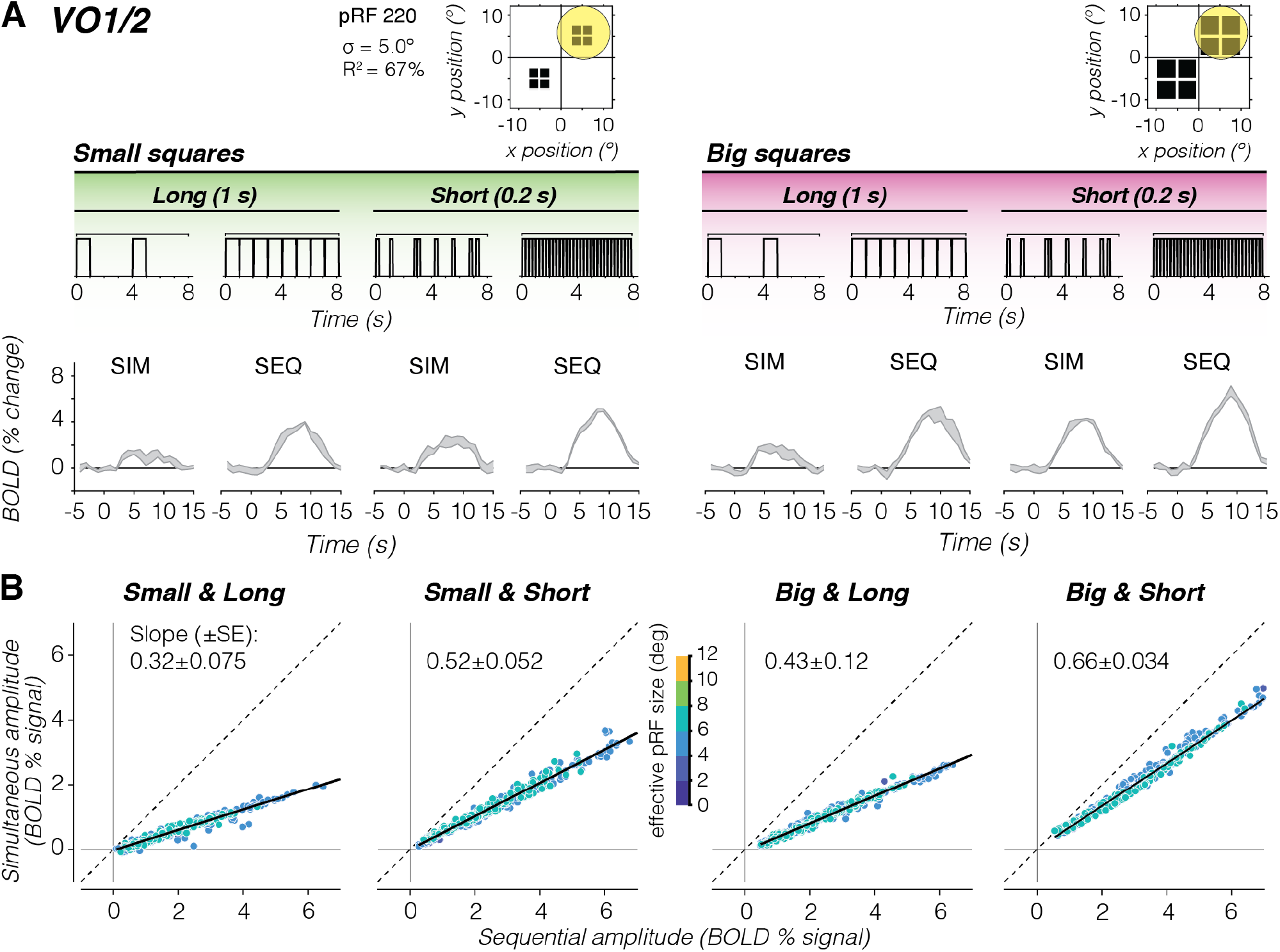
Individual VO1/2 voxels with large pRFs show strong simultaneous suppression effects. Same layout as Fig 2, but for higher-level visual area VO1/2. Data are from participant S3. **(A) Example time series of a VO1 voxel.** Voxel has a large pRF that covers all the four squares of both sizes (yellow circle). **(B) Simultaneous vs. sequential BOLD amplitude for all voxels in VO1/2.** *Dashed line:* No suppression; *Solid black line:* LMM line fit for this participant’s VO1/2 data. Slope (±SE) across voxels are indicated in each panel.

We observe this pattern of results across VO voxels. Plotting the average amplitude for simultaneous vs sequential presentations, we find a linear relationship between responses to simultaneous and sequential pairings, where voxels show simultaneous suppression and the level of suppression varies across experimental conditions (**Fig 3B**, example participant; **Supplementary Fig 1**, all participants). This relationship is not a given, as simultaneous suppression could have tapered off with response level. Instead, our data suggests that suppression can be summarized with a single slope per visual area and experimental condition.

Quantitative analyses using a LMM (R^2^=97%) revealed significant simultaneous suppression varying with stimulus size and timing, with the following suppression levels: small and long squares: 0.40±0.075 (CI_95%_=0.15–0.65), small and short squares: 0.65±0.052 (CI_95%_=0.55–0.75), big and long squares: 0.62±0.11 (CI_95%_=0.31–0.93), and big and short squares: 0.70±0.033 (CI_95%_=0.54–0.87). Notably, for stimuli of the same duration, there is larger suppression (smaller slopes) for the small vs big squares. However, for the same square size, there is larger suppression for long vs short presentation timings. This suggests that in VO1/2, in addition to the stimulus’ spatial overlap with the pRF, timing also contributes to simultaneous suppression.

### 3.3 Simultaneous suppression increases up the visual hierarchy and depends on stimulus size and presentation timing

We next quantified the relationship between responses in simultaneous vs sequential presentations across the visual hierarchy. Our data show four findings. First, in each visual area and stimulus condition, we find a linear relationship between voxels’ responses to simultaneous and sequential stimuli (**Fig 4A**, big and short stimuli; **Supplementary Fig 1**, all conditions). Second, when quantifying this linear relationship by its slope, we find that simultaneous suppression is prevalent at the voxel level in almost every visual area across participants. Third, across all stimulus conditions, we find that suppression levels progressively increase from early visual areas (V1 to V2 to V3) to intermediate areas (hV4, LO1/2, V3A/B), with the strongest simultaneous suppression in TO1/2 (**Fig 4B** and **Supplementary Table 1**). Fourth, up the visual hierarchy, simultaneous suppression levels depend on stimulus condition. In particular, higher-level visual areas show stronger suppression for long vs short presentation timings, and stronger suppression for small vs big square sizes. A two-way repeated measures ANOVA revealed significant effects of visual area (*F*(8)=23, *p*=7.3×10^-27^) and stimulus condition (*F*(3)=27, *p*=2.3×10^-15^) on suppression slopes. There was no significant interaction between stimulus condition and visual area (**Supplementary Table 2;** post-hoc Bonferroni-corrected t-tests).

**Figure 4.**
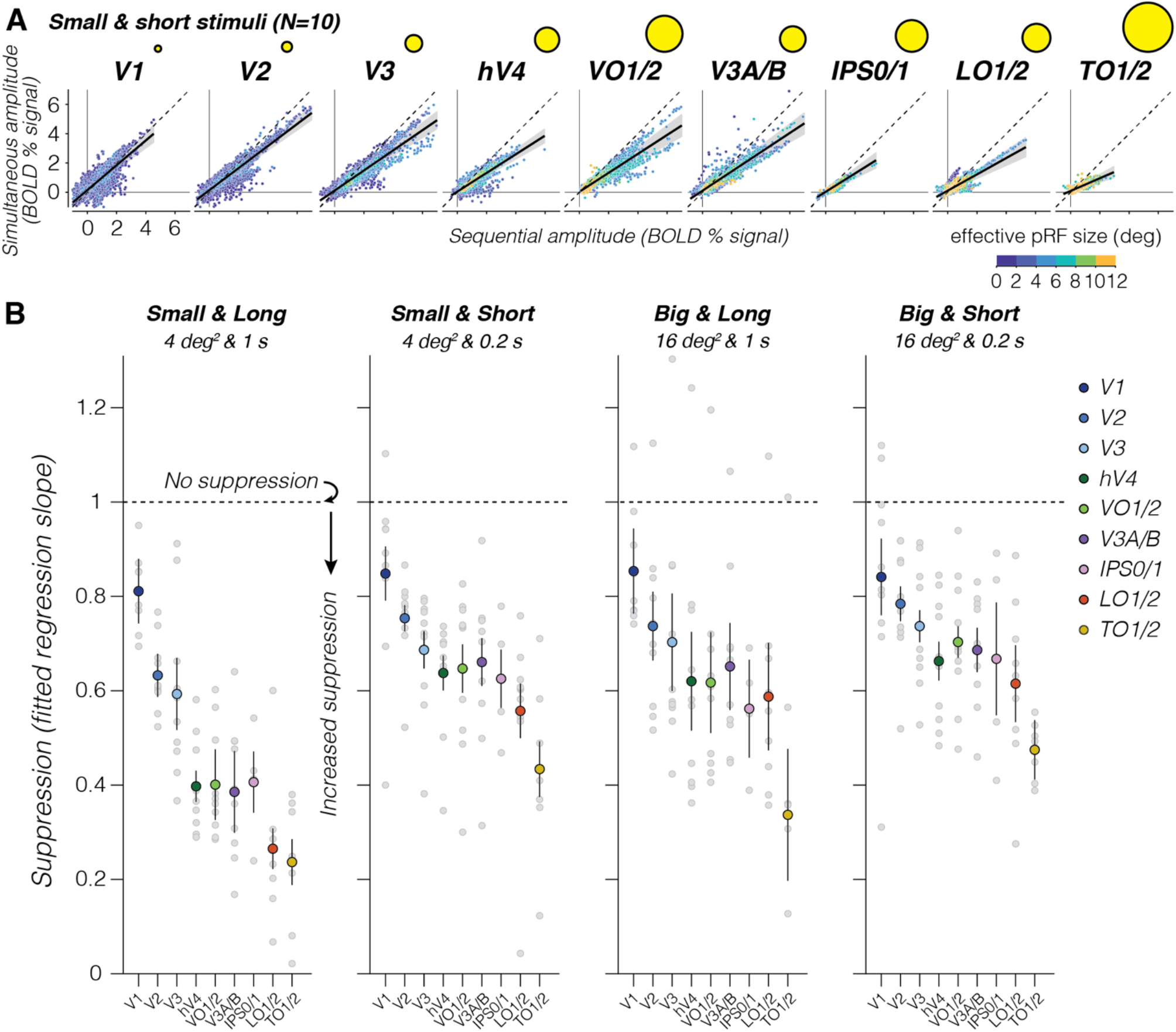
Simultaneous suppression increases up the visual hierarchy. **(A) Average sequential vs simultaneous BOLD amplitude of individual voxels for small and short stimulus condition.** Each point is a voxel, colored by effective pRF size estimated from the retinotopy data. Each panel shows data of all 10 participants. *Black solid line:* LMM fit (average across participants). *Dashed line*: identity line. *Shaded area:* CI_95%_ across participants. *Yellow circles:* illustration of average effective pRF size per area, ranging from 1° in V1 to 7.8° in TO1/2. **(B) Suppression levels for each stimulus condition and visual area.** Slopes are derived from LMM fit to simultaneous vs sequential average BOLD amplitude data from all 10 participants, for each visual area. A slope of 1 indicates no suppression. Smaller slopes indicate larger suppression. *Large colored dots:* Group average of a visual area. *Error bars*: SEM across participants. *Light gray dots:* Individual participant slopes (random effects). Early visual areas are in blue colors (V1: indigo. V2: dark blue. V3: light blue), ventral visual areas in green colors (hV4: dark green. VO1/2: light green), dorsal visual areas are in purple colors (V3A/B: purple. IPS0/1: pink), and lateral visual areas are in warm colors (LO1/2: red. TO1/2: yellow).

The increasing suppression levels across the visual hierarchy are in line with our prediction that simultaneous suppression will be stronger in visual areas that have larger pRF sizes. This relationship is evident at the level of entire visual areas (**Fig 4B**), but not across voxels within an area (**Fig 4A**). Within an area, we find similar suppression levels for voxels with pRFs that drastically vary in size (e.g., VO1/2), yet their level of suppression is predicted by a single line. Thus, while pRF size is an important predictor of simultaneous suppression at the level of an entire visual area, our data suggest that by itself, summation within pRFs that vary in size is insufficient to explain different suppression levels observed across stimulus conditions. Together, these results reveal robust simultaneous suppression at the individual voxel level that depends both on pRF size alongside stimulus size and timing parameters.

To understand how much of the observed suppression in higher-level visual areas is accumulated from earlier visual areas, we compare suppression slopes between pairs of consecutive visual areas within a processing stream. One possibility is that suppression monotonically accumulates up the visual hierarchy irrespective of stimulus condition (e.g., a consistent difference in slopes between consecutive areas). Alternatively, suppression may increase until a certain processing stage and then plateau (e.g., when the average pRF size within a visual area is large enough to encompass all square stimuli). Contrary to these predictions, we find that the difference in suppression levels between consecutive visual areas varies by stimulus condition, and suppression levels do not increase consistently across the visual hierarchy nor plateau (**Supplementary Fig 2A**). Additionally, observed differences in suppression levels between consecutive visual areas do not show a clear relationship with differences in pRF size (**Supplementary Fig 2B)** or differences in spatiotemporal compression within pRFs (**Supplementary Fig 2C**). These results suggest that there is some accumulation of simultaneous suppression up the visual hierarchy, but that accumulation alone cannot fully explain the observed simultaneous suppression levels in higher visual areas.

### 3.4 A spatiotemporal pRF modeling framework to predict simultaneous suppression at the single voxel level

To gain insight into the stimulus-driven computations that give rise to different levels of simultaneous suppression at the voxel level, we developed a computational framework that predicts the neural population response in each voxel from its pRF given the frame-by-frame stimulus sequence of the SEQ-SIM experiment (**Fig 5**). To capture the brief nature of the stimuli and the neural response, both stimulus sequence and predicted pRF responses have millisecond resolution. This neural pRF response is then convolved with the hemodynamic response function (HRF) to predict the voxel’s BOLD response and downsampled to 1 s resolution to match the fMRI acquisition (**Fig 5A**). Crucially, for each voxel, we use a single pRF model and the stimulus sequence of the entire SEQ-SIM experiment to predict its time series across all stimulus conditions at once. For all tested pRF models, spatial parameters of each voxel’s pRF are identical and estimated from the independent retinotopy experiment (**Fig 1D**).

**Figure 5.**
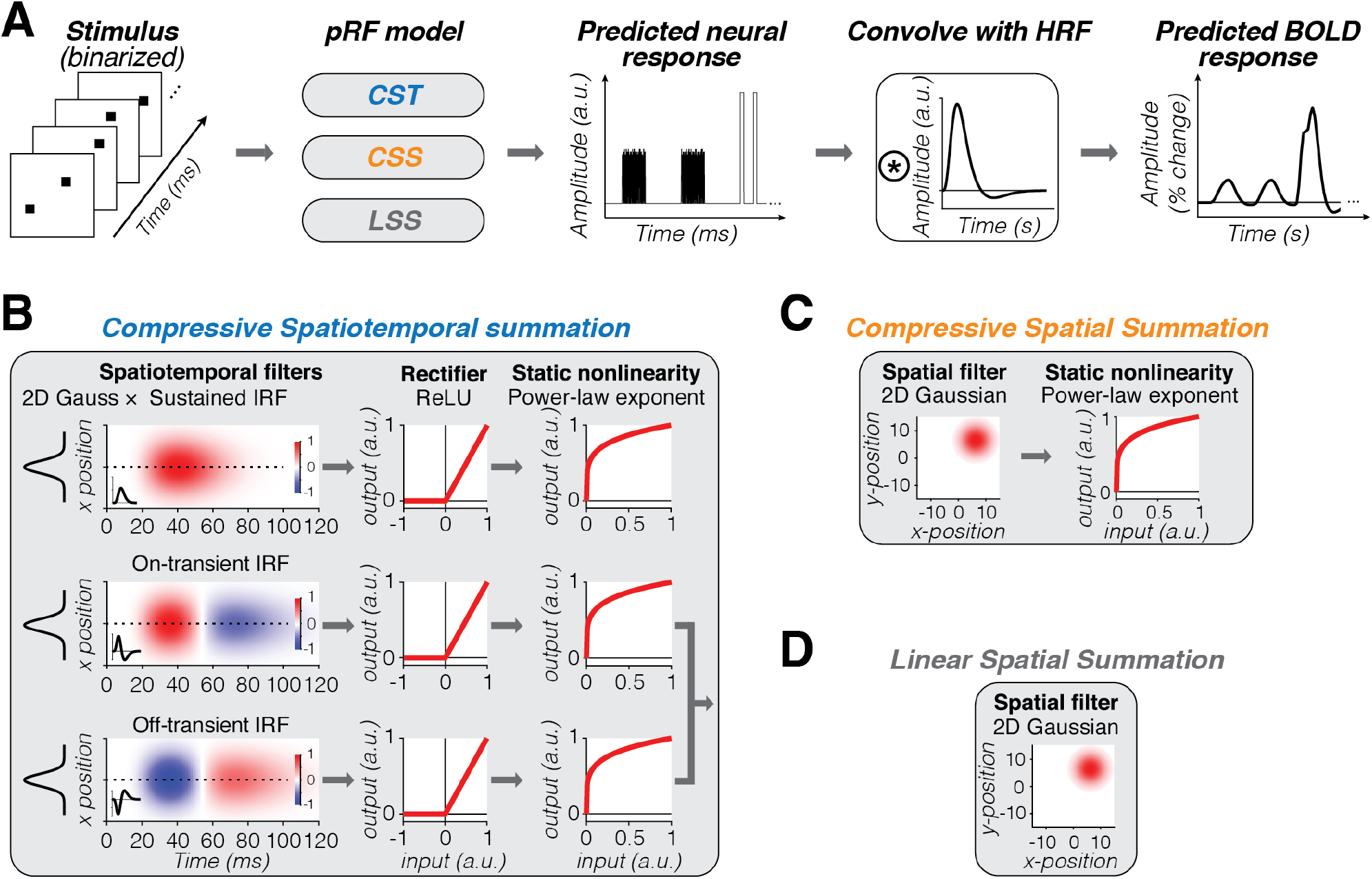
Computational modeling framework. **(A) Model overview.** *From left to right:* Given a binarized stimulus sequence and pRF model, the neural response is predicted at millisecond time resolution. This neural response is convolved with hemodynamic response function (HRF) to predict the BOLD response. After the convolution with the HRF, data are downsampled to 1-s temporal resolution (TR in SEQ-SIM experiment). **(B-D) Tested pRF models.** For each voxel, spatial pRF parameters are identical for all models and estimated from the retinotopy experiment (Fig 1D). Both CSS and LSS models sum linearly over time. For simulated pRF model predictions, see Supplementary Fig 3. **(B) Compressive Spatiotemporal summation (CST)**^51^. Temporal pRF parameters are default parameters from Stigliani *et al.*^41^. Static power-law exponent parameter (<1) is the same for all three spatiotemporal channels and fitted to each voxel’s SEQ-SIM data. The overall predicted BOLD response by the CST model is the weighted sum of the sustained and combined transient channel. **(C) Compressive spatial summation (CSS)**^27^. 2D Gaussian followed by a static compressive nonlinearity (exponent <1, estimated from retinotopy data). **(D) Linear spatial summation (LSS)**^18^. LSS pRFs sum linearly across space and time by computing the dot product between the binarized stimulus frame and the 2D Gaussian pRF.

We test three pRF models. First, a compressive spatiotemporal pRF model (CST^51^) (**Fig 5B**) to quantitatively examine if compressive spatiotemporal summation within pRFs can predict simultaneous suppression across all stimulus manipulations. The CST pRF model contains three spatiotemporal channels that have the same spatial pRF (2D Gaussian) but different neural temporal impulse response functions (IRFs): a sustained, on-transient, and off-transient channel that captures stimulus duration, onsets, and offsets; neural IRFs use default temporal pRF parameters from Stigliani *et al.*^41^. These spatiotemporal filter outputs are rectified and subjected to a compressive static nonlinearity, which produces subadditive spatiotemporal summation for both sustained and transient channels.

Second, we implement a compressive spatial summation pRF model (CSS^27^) (**Fig 5C**) to quantitatively test if subadditive spatial summation alone can explain simultaneous suppression. The CSS model has a 2D Gaussian followed by a compressive static nonlinearity and is successful in predicting spatial subadditivity in voxels with larger pRFs beyond V1.

Third, we implement a linear spatial summation pRF model (LSS^18^) (**Fig 5D**) to quantitatively test if small voxels that show little to no simultaneous suppression, such as those in V1, can be predicted by linear summation in space and time. The LSS pRF contains a 2D Gaussian for each voxel and sums stimulus input linearly over time and space. This model was also used to validate our experimental design as it predicts that, irrespective of pRF size, linear summation of stimuli in paired SEQ-SIM conditions should result in the same response, i.e., no simultaneous suppression.

We test these pRF models for four main reasons. First, they describe a neural mechanism with a receptive field restricted to part of the visual field; this restriction is needed to test the impact of stimulus location and size. Second, the identical spatial pRF across models and similar static nonlinearity implementation for compressive models (CST and CSS) allow for informative comparisons between models. Third, both compressive models have the potential to predict simultaneous suppression within this stimulus regime. Fourth, CST and CSS models have been successful in providing a comprehensive explanation for subadditive visually-driven responses across visual cortex^27,51^.

### 3.5 Comparing pRF model performance in predicting observed SEQ-SIM data

For each voxel, we generate three predicted BOLD responses, one for each tested pRF model (CST, CSS, LSS; see **Supplementary Fig 3** for example pRF model predictions). We fit each model using split-half cross-validation and quantified the cross-validated variance explained (cv-R^2^) for each voxel. This provides a principled and unbiased way to test the hypotheses.

For our example, small V1 pRF, both spatial models (LSS and CSS) predict the same BOLD response for sequential and simultaneous pairs (**Fig 6A**, bottom and middle rows). This is because the pRF covers only one small square, and consequently, spatial summation is identical across SIM and SEQ presentations. Comparing predictions to data, both LSS and CSS models capture the voxel’s response to long stimulus conditions, but underpredict the voxel’s response for short stimulus conditions, resulting in the same cv-R^2^ of 43% for this V1 voxel. In comparison, the CST pRF model best captures the response pattern across all stimulus conditions (cv-R^2^=52%), predicting no suppression and larger BOLD amplitudes for short than long stimulus conditions (**Fig 6A**, top row).

**Figure 6.**
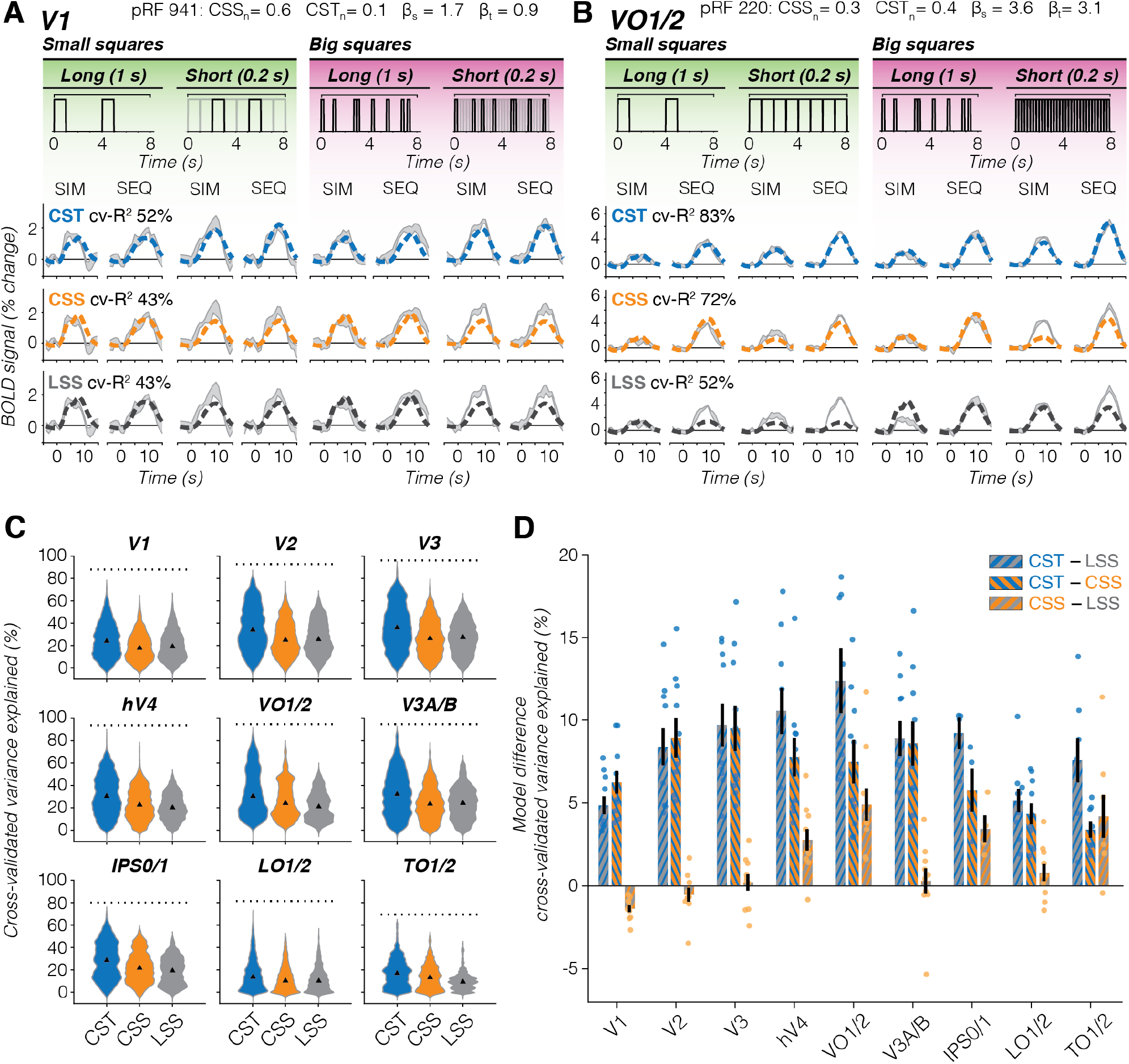
Comparison of pRF model performance. **(A) V1 example voxel.** *Gray shaded area:* Average ± SEM voxel time series. Data are from the same voxel as in Fig 2A repeated for each row. PRF model fits are shown in dashed lines. Split-half cross-validated variance explained (cv-R^2^) is computed by fitting the predicted time series to the average of odd runs and applying the modelfit to the average of even runs and vice versa. *Blue:* Compressive spatiotemporal summation model (CST, top row). *Orange:* Compressive spatial summation model (CSS, middle row). *Black:* Linear spatial summation model (LSS, bottom row). **(B) VO1/2 example voxel.** Data are from the same voxel as in Fig 3A repeated for each row. Same color scheme as panel A. **(C) Distribution of voxel-level cross-validated variance explained for each pRF model, all 10 participants.** *Triangle:* median. *Dotted line:* noise ceiling computed from max split-half reliability across participants. *Blue:* CST. *Orange*: CSS. *Gray:* LSS. Since number of voxels vary per participant and visual area, we assure equal contribution of each participant by resampling data 1000x of each participant’s visual area. **(D) Pairwise model comparison for each visual area.** *Bars:* Average across participants of the voxelwise difference in cv-R^2^ between two pRF models. *Error bars:* SEM across participants. *Individual dots:* Average difference for each participant. *Blue-gray:* CST vs LSS. *Blue-orange:* CST vs CSS. *Orange-gray:* CSS vs LSS.

When pRFs are large and cover multiple stimuli, like the example VO1/2 voxel, the LSS pRF model predicts larger responses for big than small squares, slightly higher responses for long than short presentations, and identical responses for sequential and simultaneous pairs. As such, it fails to predict the observed simultaneous suppression in all conditions (**Fig 6B**, bottom row). On the other hand, the CSS pRF model predicts simultaneous suppression because of spatial subadditivity, as well as a modest increase in response with stimulus size (**Fig 6B**, middle row). Like the LSS model, the CSS model predicts slightly larger responses for the long than short presentations of a given sequence type (SIM/SEQ). Consequently, the CSS model predicts simultaneous suppression well for the long presentations across stimulus sizes, but overpredicts simultaneous suppression for short presentations. In contrast, the CST pRF model best predicts all stimulus conditions for this example voxel: it shows simultaneous suppression, slightly larger responses for big vs small stimulus sizes, and larger responses for short vs long presentation timings (**Fig 6B**, top row).

Across all voxels and visual areas, we find that the CST pRF model best predicts our data (**Fig 6C,D**). The CST model explains more cv-R^2^ than LSS and CSS pRF models and approaches the noise ceiling in V3 and higher-level visual areas (**Fig 6C**, dotted line). A two-way repeated measures ANOVA revealed significant effects of pRF model (*F*(2)=2.6×10^3^, *p*<10^-209^) and region of interest (ROI) (*F*(8)=3.4×10^3^, *p*<10^-209^) on cv-R^2^, as well as a significant interaction between pRF model and ROI (*F*(2,8)=65, *p*=2.8×10^-209^) (post-hoc Bonferroni-corrected t-tests are reported in **Supplementary Table 3**). On average, the increase in cv-R^2^ for the CST model compared to the other models ranges from ∼5% in V1 to ∼12% in VO1/2 (**Fig 6D**). Beyond early visual cortex, the CSS model outperforms LSS, but in V1 the LSS model slightly (+1.4%) and significantly (*p*=2.7×10^-8^) explains more cv-R^2^ than the CSS model. These results suggest that V1 voxels largely sum linearly in space, but nonlinearly in time. However, across the visual hierarchy, compressive spatiotemporal summation provides a more comprehensive explanation of the empirical data.

### 3.6 To what extent do pRF models predict simultaneous suppression across visual cortex and across stimulus conditions?

To understand the underlying neural computations that generate simultaneous suppression, we use pRF models to predict the level of simultaneous suppression in each voxel and condition of the SEQ-SIM experiment. Then, we compare the model-based simultaneous suppression against the observed suppression (**Fig 7**, shaded gray bars).

**Figure 7.**
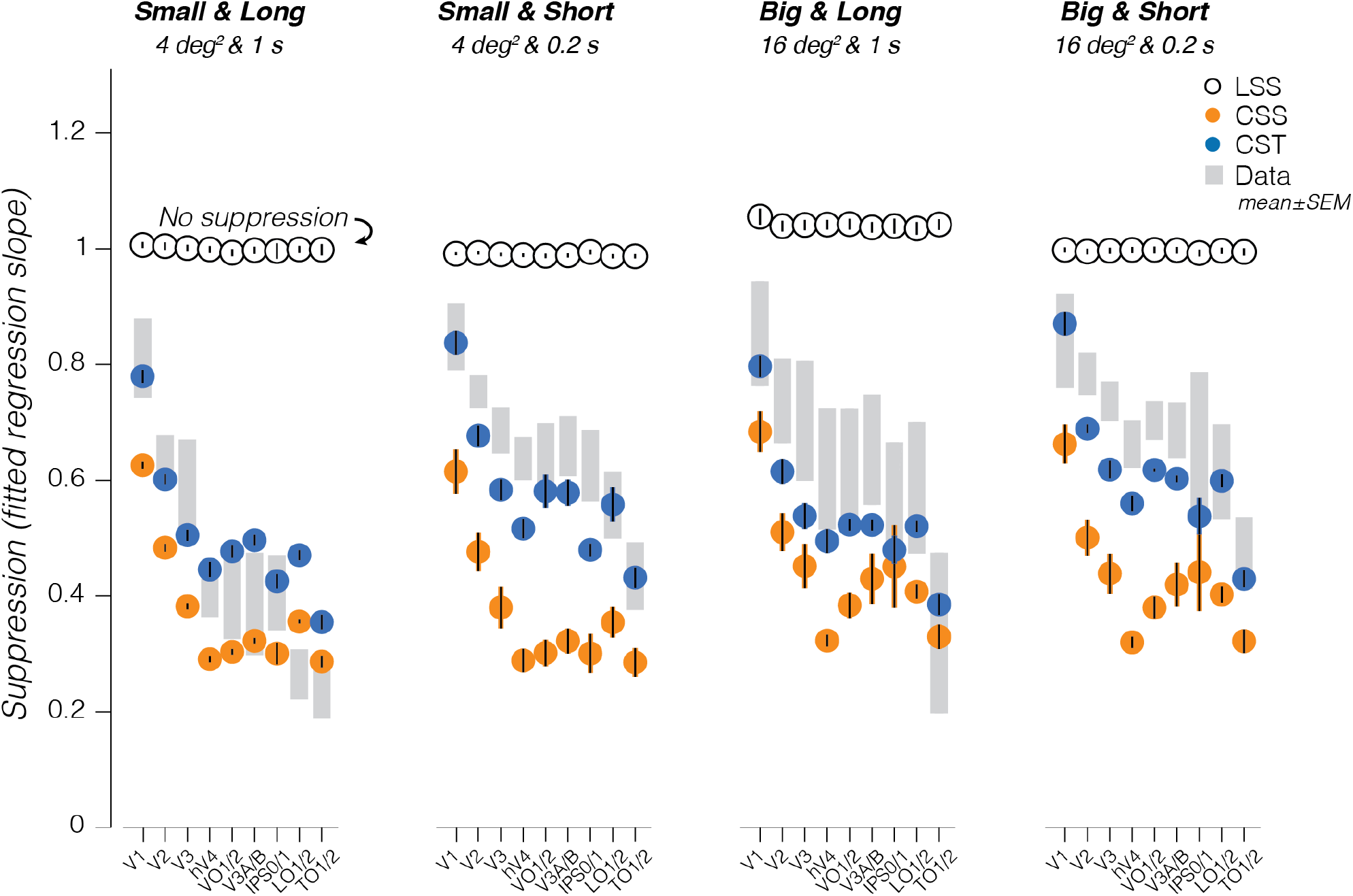
Model-based prediction of simultaneous suppression vs observed simultaneous suppresion. *Shaded gray bars:* Observed suppression levels in data, mean ± SEM across participants (same as Fig 4B). *Black open circles*: Linear spatial summation (LSS) pRF model; *Orange filled circles:* Compressive spatial summation (CSS) pRF model; *Blue filled circles:* Compressive spatiotemporal (CST) summation. Model-based points and errorbars show average and SEM across all 10 participants.

The CST model best captures simultaneous suppression across visual areas and stimulus conditions as its predictions are largely within the range of data variability (**Fig 7**, compare blue circles to shaded gray bars). Specifically, the CST model predicts (i) progressively increasing simultaneous suppression across visual hierarchy, (ii) stronger suppression for longer than shorter presentation timings for squares of the same size, and (iii) weaker suppression for bigger than smaller squares of the same timing. The CST model performs similarly with pRF parameters that are optimized using the independent spatiotemporal retinotopy experiment^51^ (**Supplementary Figs 8-9**).

The CSS model captures the progressively stronger simultaneous suppression across visual hierarchy and the observed simultaneous suppression for the long stimuli in a few visual areas (V3A/B, IPS0/1, and TO1/2), but fails to predict suppression for short stimuli and generally overpredicts the level of suppression (**Fig 7**, orange circles). In other words, the CSS model predicts much stronger simultaneous suppression levels than observed, as model predictions are consistently below the data. This overprediction is largest for short presentation timings in early (V1-V3) and ventral visual areas (hV4 and VO1). One reason for this mismodeling error is that the CSS model does not encode visual transients: it predicts stronger simultaneous suppression for small than big sizes but predicts similar simultaneous suppression for long and short presentations of the same square size.

Finally, and as expected, the LSS model does not predict simultaneous suppression altogether. This is because the LSS model sums visual inputs linearly in space and time, and we designed our experiment such that each square is shown for the same duration and location in sequential and simultaneous conditions. Therefore, the LSS model predicts the same responses for sequential and simultaneous stimulus pairings and consequently no suppression (**Fig 7**, black open circles). For the big and long squares, the LSS model predicts slightly higher responses for simultaneous vs sequential presentations. We attribute this to our experimental design, which has different inter-stimulus-intervals of individual squares between sequential and simultaneous blocks, see *Methods – LSS pRF model*).

Thus far, we primarily focused on two compressive pRF models (CST and CSS) and find that CST pRFs best predict simultaneous suppression. However, this does not rule out the possibility that other types of subadditive mechanisms within pRFs can account for simultaneous suppression. One such mechanism is center-surround suppression, which is most prevalent in early stages of visual processing (retina^57^, LGN^24^, and V1^58^). To test this possibility, we simulated Difference of Gaussians (DoG) pRFs for voxels in V1 through hV4. In these visual areas, DoG pRFs have a positive center that encompasses one or few squares while the larger, negative surround covers more squares, which may suppress the overall response in simultaneous conditions (**Supplementary Fig 7A**). Yet, we find that DoG pRFs predict no simultaneous suppression in V1 through hV4 for our stimuli (**Supplementary Fig 7B**). Instead, like the LSS model, DoG pRFs sum linearly over the spatial and temporal extent of the stimulus, resulting in similar responses across SIM and SEQ conditions.

A second potential subadditive mechanism is delayed normalization^42,44^. To test this mechanism, we implement a delayed normalization spatiotemporal (DN-ST) pRF model for 7 participants who completed a separate spatiotemporal pRF retinotopy experiment^51^. Like the CST model, the DN-ST model captures the increase in simultaneous suppression across the visual hierarchy, as well as differences in suppression levels with stimulus size and timing (**Supplementary Fig 8A**). A two-way repeated measures ANOVA reveals significant effects of pRF model (*F*(2)=1.1×10^2^, *p*=6.3×10^-47^) and ROI (*F*(7)=1.3×10^3^, *p*<10^-47^) on cv-R^2^ (**Supplementary Fig 8B**), where the DN-ST model with optimized parameters predict overall less cv-R^2^ than either CST model: 3.9% less cv-R^2^ than CST pRFs with fixed temporal parameters and 1.5% less cv-R^2^ than CST pRFs with optimized parameters. In particular, DN-ST pRFs tend to underpredict the level of simultaneous suppression for short stimulus timings (0.2 s) (**Supplementary Fig 9A**). The ANOVA also indicates a significant interaction between pRF model and ROI (*F*(2,7)=5.0, *p*=1.6×10^-9^), where both CST models perform significantly better than the DN-ST model in almost all visual areas, and the main CST pRF model explains slightly but significantly more cv-R^2^ than the optimized CST pRF model in visual areas V1, hV4, V3A/B, LO1/2, and TO1/2 (**Supplementary Table 4**, post-hoc Bonferroni-corrected t-tests).

Together, these model comparisons suggest that accounting for spatiotemporal nonlinearities rather than just spatial nonlinearities is necessary for predicting simultaneous suppression across a variety of spatial and temporal stimulus conditions.

### 3.7 What intrinsic pRF components drive the observed simultaneous suppression?

To elucidate what pRFs components predict the varying levels of simultaneous suppression across the visual system, we examine the relationship between the average suppression level and CST pRF model parameters. We find that simultaneous suppression increases with pRF size, spatiotemporal compression (*CST_n_*), and necessitates contributions from both sustained and transient temporal channels (**Fig 8**). Visual areas with larger pRF sizes tend to show stronger simultaneous suppression levels (smaller slopes, Pearson’s correlation *r*=-0.72, CI_95%_=-0.81–0.59, *p*<0.0001) (**Fig 8A**). Likewise, visual areas with stronger spatiotemporal compression within pRFs (smaller exponents) are linked to stronger simultaneous suppression levels (Pearson’s *r*=0.65, CI_95%_=0.50–0.76, *p*<0.0001) (**Fig 8B**).

**Figure 8.**
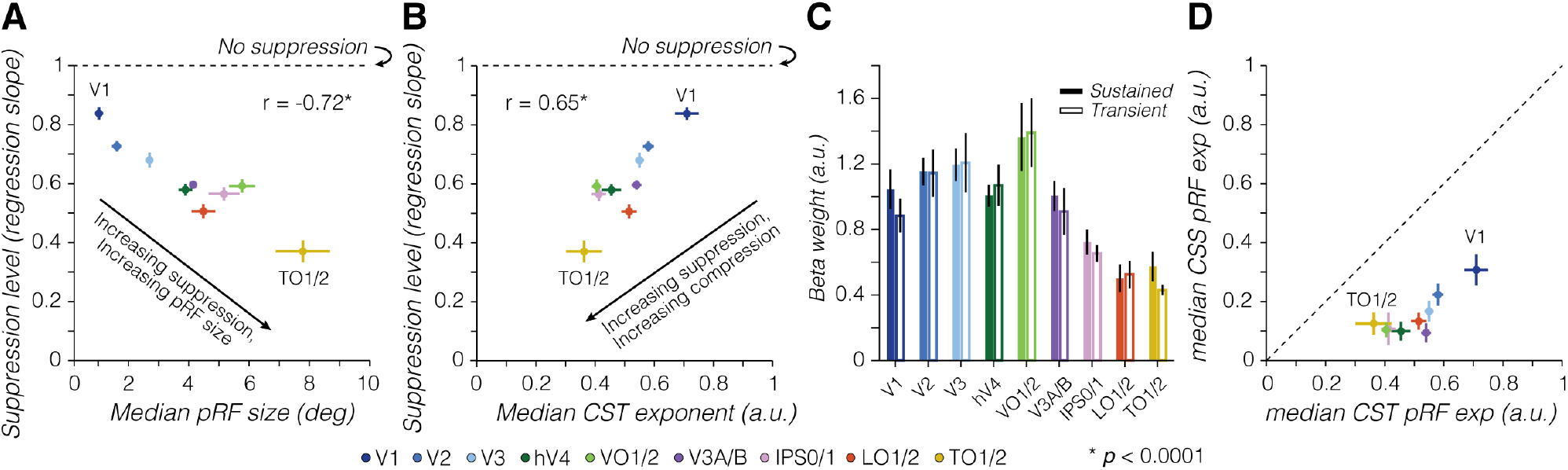
Simultaneous suppression depends on pRF size, compressive exponent, and contributions from both sustained and transient channels. In all panels: *dots/bars* show average across 10 participants. *Error bars:* SEM across 10 participants. **(A) Simultaneous suppression level vs median pRF size. (B) Simultaneous suppression level vs median CST pRF exponent.** For effective size and exponent pRF parameters, we first computed the median across pRFs of a visual area for each participant, as compressive exponent values in V1 and V2 voxels are not normally distributed (see Supplementary Fig 4), then we calculated the average median value across participants. Pearson’s correlation (*r*) is computed using individual participant data. Significant *p*-values are indicated with * (*p*<0.0001). **(C) Average β-weights of sustained and transient channels in CST pRF model.** Beta weights are averaged first within a participant’s visual area, to then average across participants per visual area. *Colored bars:* Sustained channel. *White bars:* Combined transient channel. Differences between sustained and transient channels are not significant. **(D) Median exponent pRF parameters for CSS vs CST model.** Dashed line indicates equality line.

Lastly, we find that across visual areas, both sustained and transient channels contribute to predicting single voxel BOLD responses, as their β-weights are similar (no significant difference in β-weights across channels) (**Fig 8C**). These results indicate that both sustained and transient channels are needed to predict simultaneous suppression across different stimulus size and timing conditions.

Examining the relationship between simultaneous suppression and optimized CST model parameters underscores our findings that larger pRF sizes, stronger compressive nonlinearity (i.e., smaller exponents), and contributions of both sustained and transient channels are important for predicting the level of simultaneous suppression across the visual hierarchy, whereas time constant parameters do not systematically co-vary with observed suppression levels (**Supplementary Fig 9B-D**). The DN-ST model shows similar a relationship as the CST model, where increased levels of simultaneous suppression are predicted by larger pRF sizes and stronger spatiotemporal compression within the pRF via larger semi-saturation constants in the denominator, smaller exponents, and increased exponential decay time constants (**Supplementary Fig 9F-I**). These results suggest that simultaneous suppression can be predicted by more than one implementation of a compressive spatiotemporal mechanism (static nonlinearity vs delayed divisive normalization).

Because the static nonlinearity in CST pRFs is applied to the output of spatiotemporal channels, the compressive nonlinearity is of spatiotemporal nature and cannot be separated into spatial and temporal dimensions. Nevertheless, we can gain insight into the different contributions of spatial versus spatiotemporal compression by comparing the exponent across the main CST and CSS pRF models. We find that across all visual areas, CSS pRFs have smaller exponents (resulting in more compression) than CST pRFs with a fixed time constant (**Fig 8D**). This overly strong compression by the CSS model likely explains its mismodeling of the short stimuli, where it predicts too much suppression (**Fig 7**). Interestingly, while CST pRFs with optimized spatiotemporal parameters have similarly small exponents like CSS pRFs, the CST pRFs are more accurate than CSS pRFs in predicting simultaneous suppression across stimulus conditions (**Supplementary Fig 9C**).

Overall, these results suggests that both spatial and temporal nonlinearities within pRFs are necessary to account for the observed simultaneous suppression, and ultimately interact.

## 4 Discussion

Simultaneous suppression is a decades-old, yet perplexing neurophysiological phenomenon: Why is the response to multiple stimuli presented simultaneously substantially lower compared to the response to the same stimuli presented sequentially? Here, we combined a new experimental design, varying stimulus size and presentation timing, with an innovative spatiotemporal pRF modeling framework to elucidate the stimulus-driven computations that may give rise to simultaneous suppression in individual voxels. Our results show that the level of simultaneous suppression depends not only on the spatial overlap between stimuli and the pRF, but also on the timing of stimuli and the number of visual transients. Furthermore, we find that compressive (subadditive) spatiotemporal computations by pRFs predict simultaneous suppression in each voxel across the visual hierarchy, and across various experimental conditions. These findings suggest that a stimulus-driven compressive spatiotemporal computation by pRFs generates simultaneous suppression and necessitate a rethinking of the neural mechanisms involved in simultaneous suppression.

### 4.1 Rethinking the neural mechanisms of simultaneous suppression

By investigating simultaneous suppression under a computational lens, measuring and predicting each voxel’s pRF response independently, we provide a mechanistic explanation on how the spatial overlap between the stimulus and pRF drives simultaneous suppression at the single voxel level. This confirms the longstanding hypothesis that the overlap between the receptive field and stimuli matters^8,9,12^. Additionally, we show that increasing simultaneous suppression up the visual hierarchy is predicted by both the progressive increase in pRF size and strength of spatiotemporal compression.

Crucially, we are able to explain a wide range of simultaneous suppression levels by stimulus-driven computations within pRFs alone, which necessitates a rethinking of the neural processing underlying simultaneous suppression. Thus, we propose a new idea that simultaneous suppression is a consequence of simple, stimulus-driven spatiotemporal computations rather than a result of stimuli competing for limited neural resources within receptive fields, and prioritized by task demands.

As our computational framework uses a stimulus-referred encoding model, it has predictive power. This allows future research to make new predictions about suppression levels for any stimulus sequence. The framework is also modular and can be expanded to computationally operationalize the effects of stimulus content, context, and task demands on simultaneous suppression.

### 4.2 Simultaneous suppression increases up the visual processing hierarchy, and depends on stimulus size and timing

Consistent with previous work^7–9,12^, our data show that simultaneous suppression increases up the visual hierarchy and is particularly strong in ventral visual areas (hV4 and VO1/2). Notably, we find that not only stimulus size and location, but also stimulus timing and number of visual transients affect the level of simultaneous suppression: for stimuli of the same size, longer timings (1 s) with fewer transients generated stronger suppression levels than shorter timings (0.2 s) with more transients.

Contrary to prior studies^8–12^,we find moderate levels of suppression in already in V1, despite its small receptive fields. This may be because of multiple differences between studies. First, differences in the level of analysis: we quantify simultaneous suppression at the voxel level vs ROI level in prior fMRI studies^8–12^. Second, differences in which pRFs are analyzed: we include all pRFs that overlap the stimuli, including small pRFs that partially overlap multiple squares, but electrophysiology studies used stimuli that completely overlap with neurons’ receptive fields^5–7^. Third, differences in stimulus timing: prior studies used a single stimulus timing (0.25 s per stimulus)^8–12^, which is similar to our short stimuli, for which we find weaker levels of simultaneous suppression.

### 4.3 Compressive spatiotemporal computations within pRFs can explain simultaneous suppression across visual cortex

To test whether compressive spatial summation or compressive spatiotemporal summation can predict simultaneous suppression across experimental conditions, we compared multiple pRF models. Overall, the CST pRF model provides a comprehensive explanation for simultaneous suppression across voxels spanning the ventral, dorsal, and lateral visual processing streams, as well as stimuli varying in size and presentation timing. This high performance of the CST model across all visual areas is not a given, as different models could have better predicted certain visual areas or processing streams.

Spatial pRF models captured some, but not all aspects of the observed simultaneous suppression. For instance, CSS pRFs are able to predict simultaneous suppression for long (1 s) but not shorter stimulus timings in several visual areas. As LSS, DoG, and CSS pRF models were developed for stimulus durations and intervals that evoke BOLD responses that approximately sum linearly in time, these models are limited because they do not account for visual transients. This assumption of temporal linearity is not only a limitation of the spatial pRF models we tested, but of any other pRF model that sums linearly over the stimulus duration, such as linear spatiotemporal pRF models^49^ or other center-surround pRFs^50^.

Likewise, we show that other mathematical forms of subadditive spatiotemporal summation, a delayed normalization spatiotemporal pRF model (DN-ST), can predict simultaneous suppression across stimulus conditions, as well as across visual areas. When inspecting pRF parameters of the tested spatiotemporal pRF models (CST and DN-ST), we find that pRF size and compressive nonlinearities that incorporate visual transients at the neural level are crucial for predicting simultaneous suppression.

While our spatiotemporal pRF models outperform spatial pRF models in predicting simultaneous suppression across stimuli size and timing, CST and DN-ST pRF models did not capture all spatiotemporal nonlinearities. For instance, for long stimuli, both CST and DN-ST models tend to overpredict suppression in early visual areas. Future research may improve CST and DN-ST model performance by optimizing parameters of both neural and hemodynamic temporal impulse response functions (IRFs) in each voxel^51^. Additionally, the estimation of some pRF model parameters is more sensitive to the experimental design and HRF variability (e.g., time constant) than others (e.g., pRF position)^51^. Therefore, we stress that it is important to consider how the experimental design may affect pRF parameter estimates and subsequent model performance.

We are not the first to consider temporal aspects of BOLD responses in models of the human visual system. Prior studies have suggested other hemodynamic^45–47^ and neural^35,39–42,48–50^ IRFs to capture BOLD temporal nonlinearities (see review^59^). Notwithstanding the success of these models, only the recent development of a compressive spatiotemporal pRF model^51^ with neural IRFs in units of visual degrees and milliseconds provided us with the opportunity to examine what subadditive spatiotemporal computations contribute to simultaneous suppression for the following reasons. First, a successful model needs to account for neural nonlinearities. We believe that the observed nonlinearities are of neural rather than hemodynamic origin, as electrocorticography and single unit recordings show that neural responses to brief visual stimuli evoke strong visual transients and are nonlinear^42^. In a recent study, we have shown that implementing such neural nonlinearities in a computational model rather than optimizing hemodynamic responses is necessary to predict BOLD temporal nonlinearities to brief stimuli as in the present study^51^. Second, to capture visual transients in rapid succession, the model requires neural IRFs with millisecond precision and a 50-200 ms response window rather than a 1-4s window as afforded by hemodynamic models^47,49^. Third, the model also requires a spatial pRF. While prior studies have modeled neural IRFs with millisecond time resolution^35,39–42^, without a spatial component these models are unable to predict differences in responses to one vs multiple stimuli covering a pRF.

### 4.4 Compressive spatiotemporal summation as a general computational mechanism in the visual system

A key insight from our study is that both increasing pRF size and stronger spatiotemporal compression contribute to increasing levels of simultaneous suppression up the visual processing hierarchy. This insight complements prior work^8,9^ which proposed that the progressive increase in receptive field size causes stronger simultaneous suppression in higher-level areas.

Increasing receptive field size and compression from early to higher-level visual areas have been interpreted as increasing summation windows that enhance invariance both in space^22,27,50,60,61^ and time^28,32,33,37,39,40,44^. This aligns with the idea that spatial and temporal compression of visual information share a similar processing strategy^40^ and suggests that compressive spatiotemporal summation may be a general computational principle in visual cortex. Moreover, our findings show that spatiotemporal receptive field models can be leveraged to gain insights about neural responses beyond processing of visual motion and dynamic information, such as predicting responses to rapidly presented stimuli varying in spatial locations, as in the present study. In addition to predicting the level of simultaneous suppression, the compressive spatiotemporal pRF model showed that ventral visual areas are highly sensitive to temporal properties of the visual input. These findings are in line with prior work showing that dynamic visual inputs affect not only motion-sensitive neurons in V1 and MT but also drive ventral visual stream areas V2, V3, hV4, and VO^36,62–68^.

What may be the role of compressive spatiotemporal summation? Little is known regarding to the role of compressive spatiotemporal summation outside of motion processing^69–72^. One possibility is that increasing compressive spatiotemporal summation generates representations that encode complex shape and motion information that unfolds over time^73^. This may be useful for binding different views of novel objects during unsupervised learning (associated with ventral stream functions^74,75^) or for perceiving complex visual dynamics, actions, and social interactions (associated with lateral stream functions^76–78^). Another possibility is that spatiotemporal compression within pRFs may enable neurons to prioritize novel visual information^13,79^. This may be beneficial for visual search^1,2^ or short-term visual working memory by converting redundant visual information into a more efficient representation^80^. However, spatiotemporal compression may also limit visual processing capacity, affecting downstream cognitive processes such as worse memory for simultaneously vs sequentially presented items^81^. Thus, an important future direction is characterizing and computationally linking the neural phenomenon of simultaneous suppression to behaviors such as visual capacity, and testing what computational mechanisms generalize across scenarios and tasks. In sum, our empirical data and voxel-wise pRF modeling approach call for a rethinking of the neural mechanisms that drive simultaneous suppression and suggest that suppression is a byproduct of compressive spatiotemporal computations. These findings provide exciting new opportunities to computationally understand how stimulus content, context, and task demands affect simultaneous suppression and visual processing capacity more broadly.

## 5 Methods

### 5.1 Participants

Ten participants (6 self-identified as female, 4 self-identified as male, ages 22-53 years, M = 30.1 years, SD = 8.7 years) with normal or corrected-to-normal vision participated in a retinotopy and SEQ-SIM fMRI experiment. Participants gave written informed consent, were compensated for their time, and all procedures were approved by the Stanford Internal Review Board on Human Subjects Research.

### 5.2 Stimuli & experimental design

Stimuli were generated using MATLAB R2017b (MathWorks, MA, USA) and PsychToolbox^82^ on an Apple MacBook Pro laptop. Images were presented using an Eiki LC-WUL100L projector (Eiki International, Inc., CA, USA) on a rear-projection screen via two large mirrors placed at the back of the MRI scanner bed. The projected image had a resolution of 1920×1080 pixels, resulting in a field-of-view of ∼38×24°, and refresh rate of 60 Hz. The display was calibrated using a linearized lookup table.

#### Retinotopy experiment

Participants completed four 3.4-minute runs, where bar stimuli cropped from colorful cartoons traversed across a 24×24° circular aperture (Toonotopy^56^). Cartoon images inside the bar changed randomly at 8 Hz. The bar swept in 12 discrete steps, 2-s per bar position, for 4 orientations (0°, 45°, 90°, 135°) and 2 motion directions for each orientation. Observers fixated on a central dot (diameter = 0.12°) and pressed a button every time the fixation dot changed color (semi-random intervals, 6–36 s). Due to a coding error, button presses were only recorded for 3 participants, who performed at ceiling (M = 98.7% correct, SD = 1.2%).

#### SEQ-SIM experiment

Participants completed eight ∼5.5-minute runs (except for participant S5, completing six runs), where 8 squares were presented sequentially or simultaneously while fixating: 4 squares in the lower right quadrant and 4 squares in the upper left quadrant. Both sequential and simultaneous conditions used two presentation timings (short: 0.2 s and long: 1 s) and two sizes (small: 2×2° and big: 4×4°), resulting in eight conditions. We did not consider sex and/or gender in this study design.

##### Stimuli

Squares were randomly cropped from colorful cartoons and placed on a mean luminance gray background. To ensure square stimuli would elicit responses in visual cortex, squares with little to no contrast were excluded (normalized root mean square contrast across pixels < 10%). The content of individual squares differed for each trial and quadrant, and never repeated within a run. Within a quadrant, squares had a 2-by-2 layout with a 0.82° gap between them, centered at ∼7.1° eccentricity ([x,y] = [5°,5°]). Both sizes used identical gap and eccentricity, such that 4 small squares extended horizontally and vertically from 2.59° to 7.41°, and big squares extended from 0.59° to 9.41°. The lower right and upper left quadrant had the same square locations but mirrored horizontally and vertically.

##### Experimental Design

Stimuli were shown in ∼8 s blocks, interspersed by 12-s blank periods. Each run started with a 6-s countdown and 12-s blank and ended with a 12-s blank. Each condition was repeated four times in a pseudo-randomized order across two runs. The block order, as well as individual square presentation within a block, differed across runs. Each participant was assigned a unique pair of runs, which were repeated four times (three for participant S5) within the experiment with different square content (see example: https://osf.io/7rqf4).

Sequential and simultaneous conditions had 8 trials per block for short stimuli and 2 trials per block for long stimuli. We used different trial-per-block ratios such that short and long conditions had a similar total block duration while the number of visual transients quadrupled (16 vs 64)—matching the increase between small and big square sizes (4 vs 16 deg^2^). In a sequential trial, the four squares in each quadrant appeared one at a time, in random order, with a 33-ms inter-stimulus-interval (ISI) between squares. In a simultaneous trial, all four squares in a quadrant appeared at once for the same duration and location followed by a mean luminance gray display to match duration of a sequential trial.

Block onsets and stimulus conditions were identical across quadrants, but timing and order of individual square appearances were independently determined per quadrant. In simultaneous blocks with long stimulus presentations, stimuli in the first trial were presented at block onset to match sequential blocks. Stimuli of the second trial were presented 4 s later to avoid 7-s gaps between stimuli within a block. In simultaneous blocks with short presentations, stimuli in the first trial were also locked to block onset, but onset of stimuli in the following 7 trials was randomized within a trial.

##### Task & Behavioral performance

Participants performed a 1-back letter RSVP task at fixation and pressed a button when a letter repeated (1/9 probability). The letters (diameter of ∼0.5°) updated at 1.5 Hz, alternating between black and white colors, and randomly drawn from a predefined list (‘A’, ‘S’, ‘D’, ‘F’, ‘G’, ‘H’, ‘J’, ‘K’, ‘B’, ‘P’). Participants had a 0.83-s response window after a letter appeared and performance was displayed after every run. Outside the scanner, participants did 1-minute practice runs until they reached at least 70% correct before starting the experiment. In the scanner, participants performed the task well (M = 88% correct, SD = 8.2%), ranging from 68–95%, and average false alarm rate of 2%. These behavioral data are confirmed by steady fixation in eye movement data (**Supplementary Fig 5**) and indicate that participants were fixating throughout experimental runs.

### 5.3 MRI data acquisition

Participant’s structural and functional data were collected using a 3T GE Signa MR750 scanner located in the Center for Cognitive and Neurobiological Imaging at Stanford University. Whole brain T1-weighted anatomy data were acquired using a BRAVO pulse sequence (1 mm^3^ isotropic, inversion time=450 ms, TE=2.912 ms, FA=12°), using a Nova 32-channel head coil. Functional data were collected using a Nova 16-channel coil, using a T2*-sensitive gradient echo planar imaging sequence (2.4 mm^3^ isotropic, FoV=192 mm, TE=30 ms, FA=62°). EPI slice prescriptions were oblique, roughly perpendicular to the calcarine sulcus. Retinotopy experiment used a TR of 2000 ms and 28 slices. SEQ-SIM experiment used a TR of 1000 ms and 14 slices. A T1-weighted inplane image (0.75×0.75×2.4 mm) was collected with the same coil and slice prescription as the functional scans to align functional and anatomical scans.

Left eye gaze data of 9 participants were continuously recorded in each SEQ-SIM run at 1000 Hz using an EyeLink 1000 (SR Research Ltd., Osgoode, ON, Canada). Eye position calibration and validation was conducted before the first run, using a 5-point grid. We could not collect eye gaze data in one participant due to constraints in the mirror setup. Four participants were excluded prior to analysis due to excessive measurement noise. Analysis details for eye gaze data are in the *Supplementary Material* above **Supplementary Fig 5**.

### 5.4 MRI data analysis

#### 5.4.1 Data availability

Data are provided with this paper and publicly available at https://osf.io/rpuhs/.

#### 5.4.2 Code availability

Analyses were conducted in MATLAB (R2020b) and FreeSurfer’s auto-segmentation^83^ (v6.0; http://surfer.nmr.mgh.harvard.edu/). Analysis code is publicly available on GitHub: https://github.com/VPNL/simseqPRF and https://github.com/VPNL/spatiotemporalPRFs.

#### 5.4.3 Preprocessing

Whole-brain T1-weighted scans were aligned to the AC-PC line using SPM12 (https://github.com/spm/spm12) and auto-segmented with FreeSurfer’s *recon-all* algorithm. Functional data were slice-time corrected, motion corrected, drift corrected, converted to percent signal change using the *Vistasoft* toolbox (https://github.com/vistalab/vistasoft). Participants’ functional scans were aligned with the inplane to their whole brain anatomy scan, using a coarse, followed by a fine 3D rigid body alignment (6 DoF) using the *alignvolumedata_auto* toolbox (https://github.com/cvnlab/alignvolumedata). The first 8 (SEQ-SIM) or 6 (Retinotopy) volumes of each functional scan were removed to avoid data with unstable magnetization.

##### Retinotopy analysis

Retinotopy runs were averaged and analyzed with Vistasoft’s compressive spatial summation pRF model (CSS)^27^ using a 2-stage optimization (coarse grid-fit, followed by fine search-fit). For each voxel, this resulted in 2D Gaussian pRF with center coordinates (*x_0_*, *y_0_*) in degrees, pRF standard deviation (*σ*) in degrees and pRF static nonlinearity exponent (CSS*_n_*) ranging from 0.01 to 1. To avoid pRFs that are not visually responsive, we selected pRFs with R^2^ ≥20% in the retinotopy experiment, similar to previous pRF publications^56,84^.

##### Defining visual areas

Spatial pRF parameters were converted to polar angle and eccentricity maps and projected to participant’s native cortical surface using nearest neighbor interpolation. Visual field maps were used to define the following visual areas: V1, V2, and V3^85^, hV4 and VO1/2^86^, LO1/2 and TO1/2^87^, and V3A/B and IPS0/1^88^.

##### Defining ROIs and selecting voxels

For each visual area, we selected voxels with pRFs centers within the circumference of the big squares in the SEQ-SIM experiment, that is, within an 8.82×8.82° square located 0.59° to 9.41° from display center in both x- and y-dimensions in each quadrant. From these voxels, we used those with corresponding data from the SEQ-SIM experiment. Overall, we obtained data in most participants’ visual areas, except 6 participants who had insufficient coverage of IPS0/1 and 2 participants who had insufficient coverage of TO1/2, due to fewer slices in the SEQ-SIM than retinotopy experiment.

##### SEQ-SIM analysis

We excluded voxels with a split-half reliability <10% to filter out those voxels with little to no visual response. Excluded voxels were mostly from V1 and V2, with small pRFs that fell in between stimuli or on the border of stimuli. The two unique SEQ-SIM runs were concatenated for each repeat. When applying split-half cross-validation for model fitting, the 4 concatenated runs were split into two odd and two even runs, and averaged within each half.

### 5.5 pRF modeling framework

Our modeling framework contained three main pRF models: (i) LSS, to test linear spatial summation^18^, (ii) CSS, to test compressive spatial summation^27^, and (iii) CST, to test compressive spatiotemporal summation^51^. Both LSS and CSS models linearly sum over the temporal duration of the stimulus.

Each model’s input is a 3D binarized stimulus sequence, pixels by pixels (in visual degrees) by time (milliseconds). Each pRF is applied to each frame of the stimulus sequence to predict the neural pRF response. For each model, this neural response is then convolved with a canonical hemodynamic response function (HRF) (double-gamma SPM default) and downsampled to the fMRI acquisition TR. This results in a predicted BOLD response for the entire stimulus sequence. For each pRF that overlapped stimuli in SEQ-SIM experiment, predictions were computed for each unique 5.5-min run, and then concatenated for the two unique runs. Importantly, concatenated runs contained all 8 stimulus conditions, requiring each model to predict all conditions simultaneously.

We model spatial pRF parameters in each voxel using independent retinotopy data and then test which type of spatial and/or spatiotemporal pRF computations predict simultaneous suppression in each voxel in the main experiment.

#### LSS pRF model

The LSS model has a circular 2D Gaussian pRF with an area summing to 1. The pRF computes the dot product between the 2D Gaussian and stimulus sequence at each time point to predict the response of the neural population within a voxel:

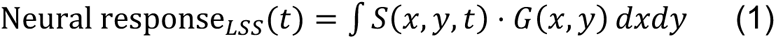

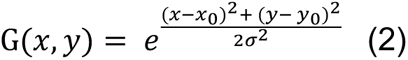

where *t* is time in ms, *S* is the stimulus sequence with visual field positions *(x, y)* in visual degrees by time in ms, and *G* is a circular 2D Gaussian centered at visual field positions *(x_0_, y_0_)* with size *σ* in visual degrees.

This model sums inputs linearly in visual space and time, and typically predicts the same BOLD response for sequential and simultaneous trials that are matched in stimulus size, location, and duration. For longer stimulus durations, the LSS model occasionally predicts larger responses for simultaneous than sequential conditions, due to a difference in ISI between the two conditions. Specifically, the randomized square onset causes sequential ISIs to range from 1–7s, which by chance can be longer than the fixed 4-s simultaneous ISI—especially for small pRFs that overlap a single square. For these scenarios, the LSS model predicts the BOLD responses accumulate less in the sequential than simultaneous block.

#### CSS pRF model

The CSS model has the same spatial pRF model as the LSS model, followed by a static power-law nonlinearity:

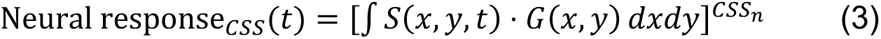

where *t* is time in ms, the power-law exponent (*CSS_n_*) is bound between 0.01–1 and therefore results in compressive (subadditive) summation.

#### CST pRF model

The CST model contains three spatiotemporal channels for each voxel. Each channel has the same spatial pRF as the LSS model, which is combined with a sustained, on-transient, or off-transient neural temporal impulse response function (IRF).

For the main analysis, the sustained, on-transient, and off-transient IRFs are identical across voxels, where neural IRFs are based on the following gamma function:

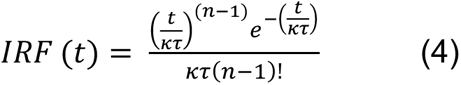

where *t* is time in ms, *κ* is the time constant ratio parameter, τ (tau) is the time constant parameter in ms, and *n* is an exponent parameter.

The sustained IRF is a monophasic gamma function as in (4) with *κ* = 1, τ = 49.3 ms and *n* = 9, resulting in a peak response between 40-50 ms. The on-transient IRF is the difference of two gamma functions: the sustained IRF and a second gamma function as in (4) with parameters: τ = 4.93 ms, *κ* = 1.33, *n* = 10, resulting in a biphasic function that generates a brief response at stimulus onset. These IRF parameters are default V1 parameters from Stigliani *et al.*^39,41^, which are based on human psychophysics^89^. The off-transient IRF is identical to the on-transient IRF but with opposite sign, generating a response at stimulus offset. The area under the sustained IRF sums to 1, and area under each transient IRF sums to 0.

For each channel, we compute the spatiotemporal pRF response to a stimulus sequence by first applying the dot product between the spatial pRF and the stimulus sequence at each time point (millisecond resolution). This spatial pRF output is then convolved with the channel’s neural temporal IRFs. The resulting spatiotemporal response is then rectified to remove negative values in the transient channels as we reasoned that either on- or off-transient responses will increase BOLD responses (sustained responses are always positive). The rectified sustained, on-transient, and off-transient channel responses are then subject to the same static power-law nonlinearity, controlled by the exponent parameter (*CST_n_*), resulting in the following neural response for each channel:

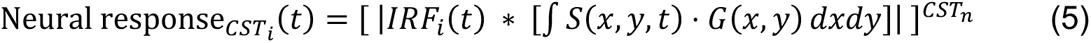

where *i = [1,2,3]* indicating the sustained, on-transient, or off-transient channel, *t* is time in ms, ∗ indicates convolution, and the power-law exponent (*CST_n_*) is bound between 0.1–1, compressing the spatiotemporal channel responses. We use a compressive nonlinearity as we reasoned that simultaneous suppression is due to subadditive summation.

After predicting neural responses for each channel, we sum on- and off-transient channels. The sustained channel and combined transient channels are then convolved with the HRF and downsampled to 1 s to predict fMRI data. The voxel’s response is the weighted sum of the two (*β_S_*, *β_T_*) time series. More details about the CST model can be found in Kim *et al.*^51^.

#### Alternative pRF models

In addition to the main pRF models, we tested two alternative spatiotemporal pRF models—CST_opt_ and DN-ST—for 7 of our participants who took part in a separate spatiotemporal retinotopy experiment^51^, and a Difference of Gaussians (DoG) pRF model^48^. We used CST_opt_ and DN-ST pRF model parameters that were optimized with a default HRF to match the main CST pRF model.

##### CST_opt_

The same model like the above-mentioned CST model, but using optimized spatial and temporal pRF parameters for each voxel, estimated from the separate Kim *et al.* experiment^51^, to understand the impact of using mostly fixed pRF parameters in the CST model. When describing CST_opt_ model performance, we refer to the main CST model as CST_fix_, because most its pRF parameters are fixed (spatial parameters from the Toonotopy experiment, a fixed time constant from Stigliani *et al.*^39,41^), except for the exponent (*CST_n_*), which is estimated from the SEQ-SIM data.

##### DN-ST

A divisive normalization spatiotemporal pRF model, to test if compressive spatiotemporal summation within the pRF with a different mathematical form can equally well predict the SEQ-SIM data. The DN-ST model implements subadditive summation using divisive normalization and an exponential decay function^40,42,44^. As for the CST_opt_, DN-ST model parameters are optimized for each voxel from a separate spatiotemporal retinotopy experiment by Kim *et al.*^51^.

#### DN-ST pRF model

The DN-ST model has a 2D circular Gaussian spatial pRF combined with a temporal IRF that contains a divisive normalization and an exponential decay function as implemented in Kim *et al*.^51^:

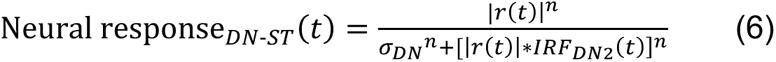

where *t* is time in ms, *σ_DN_* is a semi-saturation constant, *n* is the exponent, and *r(t)* is the linear component of the neural response (computed as the convolution between the neural temporal impulse response function *IRF_DN1_* and the spatial pRF response to the stimulus sequence (same as equation 1):

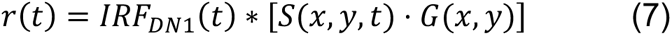

where *IRF_DN1_* is a gamma function with time *t* in ms and first time constant τ_1_ in ms:

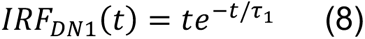

The second temporal IRF in the denominator of the neural response acts as a low-pass filter using an exponential decay function:

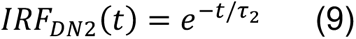

where *t* is time in ms and τ_2_ is the second time constant parameter in ms. More details about the DN-ST model, spatiotemporal retinotopy experiment, and optimization procedure can be found in Kim *et al.*^51^.

#### Difference of Gaussians (DoG) pRF model

To test whether center-surround DoG pRFs predict simultaneous suppression, we simulate each voxel’s pRF as two 2D Gaussians, a center from which a larger surround is subtracted^48,50^. We simulate DoG pRFs only for V1 to hV4 for three reasons. First, the surround pRF needs to encompass more visual input than the center pRF to predict simultaneous suppression. These visual areas have pRFs where the smaller pRF center would encompass a subset of square stimuli and the larger suppressive surround pRF would encompass more squares than the center. Second, the surround pRF of voxels in visual areas beyond hV4 would likely extend far beyond the visual display, unlikely to make a significant contribution to the response. Third, the effects of surround suppression are most prominent at early processing stages, including LGN^24^ and early visual cortex (V1-V3^48,50^).

The center Gaussian is identical to the LSS pRF, estimated from the retinotopy session (equation 1). The surround Gaussian has the same center position with a larger size, where the scale factor was based on the average center/surround size ratio from Aqil *et al.*^50^; V1: 7.4, V2: 6.8, V3: 7.3, and hV4: 5.8 times the center size. We used a constant scaling for all voxels within the same visual area, because directly estimating DoG pRFs from the independent retinotopy data using the approach by Zuiderbaan *et al.*^48^ resulted in unstable model fits. This instability is likely due to the relatively few and short blank periods in our retinotopy experiment compared to Zuiderbaan *et al.*, which has 4 x 30s-blank periods for each 5.5-min run.

#### Fixed and optimized pRF parameters

Spatial pRF parameters were independently estimated from each participant’s retinotopy experiment using the CSS pRF model, resulting in a pRF center (*x_0_, y_0_*), standard deviation (*σ*) and exponent (*CSS_n_*) parameter for each voxel. The standard deviation and exponent parameter trade-off in the CSS model (see Kay *et al.*^27^), where 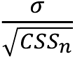 approximates the effective pRF size: the standard deviation (*σ*) estimated with a linear pRF model (LSS, no spatial compression). Therefore, to reconstruct CSS pRFs, we use each voxel’s estimated CSS parameters (*x_0_, y_0_*, *σ,* and *CSS_n_*). To reconstruct LSS and CST pRFs, we use the same estimated pRF center (*x_0_, y_0_*), but for the standard deviation (*σ*) we use the effective pRF size 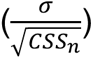.

The time constant parameter (τ) for the temporal IRF in the main CST model is fixed (from Stigliani *et al.*^39,41^) and we only optimized the exponent (*CST_n_*) using a grid-fit approach for each voxel. The best fitting *CST_n_* was determined by systematically evaluating goodness-of-fit of predicted time series with *CST_n_* between 0.1–1 (0.05 steps) and selecting the *CST_n_* resulting in the highest cross-validated R^2^. We used a grid-fit instead of a search-fit optimization approach to avoid estimates getting stuck in a local minimum.

We use a fixed temporal IRF for the following reasons. First, we predicted that the main driver of the suppression effect will be the compressive static nonlinearity (power-law exponent). Second, by using the same spatial parameters for all pRF models estimated from the independent retinotopy experiment, the model comparison will be more informative as differences in model performance are due to differences in nonlinear computations, not spatial position. Third, to estimate all CST parameters we need a separate spatiotemporal pRF retinotopy experiment^51^. We have such data for 7 out of 10 participants, which we used for comparing CST_fix_, CST_opt_, and DN-ST spatiotemporal models. For this comparison, we restricted our analysis to voxels whose pRF centers overlap the square stimuli in the SEQ-SIM experiment, as well as voxels whose variance explained by the spatiotemporal pRF model was 20% or higher, and whose split-half reliability in the SEQ-SIM experiment was 10% or higher. As in the spatiotemporal pRF retinotopy experiment, we excluded voxels with CST time constants (tau) larger than 1000 ms. This sub-selection of voxels results in a substantially smaller number of voxels per visual area (∼60%) than we used to compare the main pRF models and resulted in the removal IPS0/1 results as only two participants contributed to this visual area.

### 5.6 Model fitting

We fitted each voxel’s pRF model prediction separately to data, using a split-half cross-validation procedure. The maximum height of predicted BOLD time series was normalized to 1 and we added a column of 1’s to capture response offset. This resulted in two regressors (*β_0_*, *β_1_*) for LSS and CSS models, and three regressors (*β_0_*, *β_S_*, *β_T_*) for CST. We used linear regression (ordinary least squares) to fit these regressors to the voxel’s observed time series, separately for odd and even splits. To determine model goodness-of-fit (variance explained), we computed the cross-validated coefficient of determination (cv-R^2^) by using the scaled predicted time series of one split to predict observed time series from the other split and vice versa (i.e., β-weights are fixed and not refitted). Cv-R^2^ values and β-weights were averaged across split halves for each voxel. Split-half reliability across runs was used as the noise ceiling.

To check whether CST model performance could be inflated by the extra regressor, we also computed cross-validated adjusted-R^2^, which penalizes goodness-of-fit for the number of time points and explanatory variables. The adjusted-R^2^ values were almost numerically identical to R^2^ and did not significantly affect our results nor statistical comparisons.

### 5.7 Linear mixed model

To quantify simultaneous suppression, we fitted a linear mixed model (LMM) to all participant’s voxels within a visual area with MATLAB’s *fitlme.m*, using the maximum likelihood fitting method. This LMM predicted the average simultaneous BOLD response of each voxel as a function of the average sequential BOLD response, for each stimulus condition (fixed interaction effect), allowing for a random intercept and slope per participant and stimulus condition (random interaction effect):

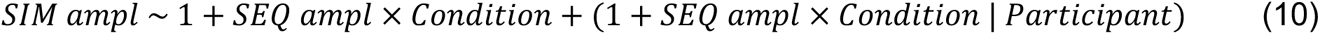

where *SIM ampl* and *SEQ ampl* are a matrix (nr voxels x 4) with continuous values, *Condition* is a categorial vector (1 x 4), and *Participant* is the group level for the random effects (10 participants for main pRF models, 7 participants for DN-ST and CST_opt_ models).

This LMM captured our data well (mean R^2^ = 90%, SD = 6.6%), with V1: 86%, V2: 94%, V3: 94%, hV4: 92%, VO1/2: 97%, V3A/B: 95%, IPS0/1: 88%, LO1/2: 85%, and TO1/2: 76% variance explained. We tested this LMM to three alternative LMMs: (i) mean sequential amplitude as a fixed factor (no condition interaction effect) with one random intercept per participant, (ii) a fixed interaction effect with a single intercept per participant, identical for each stimulus condition, and (iii) a fixed interaction effect with a random participant intercept for each condition. Despite having more degrees of freedom (45) than the alternative LMMs (4, 10, and 19), the main LMM was a better fit to the data as it had a significantly higher log-likelihood than alternative LMMs, and lower AIC and BIC for each visual area (F-test *p*<0.00001) (**Supplementary Fig 6**).

### 5.8 Summarizing results

#### BOLD time series

Both observed and predicted run time series were averaged across split-halves and segmented into 23-TR time windows. These time windows spanned from 4 s pre-block onset, 8 s stimulus block, to 11 s post-block. For each voxel, we took the average time window and standard error of the mean (SEM) across 4 repeats.

#### Seq vs Sim BOLD amplitude

The average data and model time windows were summarized into 8 values per voxel (one per condition), by averaging the BOLD response within a 9-TR window centered on the peak, spanning from either 4–12s or 5–13s after stimulus block onset. These values were used in LMMs and scatter plots. We used a variable start per condition and visual area because the BOLD accumulation rate differed. The start was determined by averaging (data or model) time windows across voxels within a visual area and condition, into a “grand mean” time window and finding the first TR after block onset where the BOLD response exceeded 10% of the total cumulative sum. This averaging window was applied to all voxels within a visual area.

#### Simultaneous suppression effects

We summarized LMM results for each condition and visual area as line fits with 95%-confidence intervals (CI_95%_) using the slope and intercept of the individual participants (**Fig 2B** and **3B**) or average across participants (**Fig 4A**). For **Fig 4B**, we summarized the simultaneous suppression level using the average slope and SEM across participants. For **Fig 8**, we first averaged slopes across conditions within a participant, and then average slopes across participants (± SEM). For **Supplementary Fig 2A**, we assessed differences in suppression levels between consecutive visual areas (suppression deltas). We computed suppression deltas separately for each participant by subtracting each participant’s regression slopes of the earlier visual area from the later visual area. Visual area pairs are chosen based on a feedforward visual hierarchy for each visual processing stream^76–78^. We then calculated the average deltas across 10 participants (± SEM) for each visual area pair and each stimulus condition. For **Supplementary Fig 2B**, we first calculated the average slopes across stimulus conditions within a participant, then computed the delta suppression slopes between pairs of visual areas within participants, and then computed the average delta suppression across participants (± SEM).

#### PRF parameters

We resampled pRF size, exponents (*CSS_n_*, *CST_n_*, DN-ST *n*), time constants (CST_opt_ tau, DN-ST tau_1_ and tau_2_), DN-ST semi-saturation constants, and CST *β_S_* and *β_T_* 1000x with replacement within a participant’s visual area, because the number of voxels varied across areas and participants. For pRF size, exponent, time constant, and semi-saturation constant, we report the median resampled parameter for each participant and visual area because the V1 and V2 *CST_n_* were not normally distributed (see **Supplementary Fig 4**). CST *β_S_* and *β_T_* were normally distributed; hence, we report the average resampled beta weights per participant and visual area. For group results, we report the average (± SEM) across participants’ mean or median resampled parameter value, for each visual area. For **Supplementary Figs 2B and 2C**, we first compute the difference in median pRF size (or CST exponent) for each visual area pair within participants, and then average delta pRF size (or CST exponent) values across participants (± SEM).

### 5.9 Statistical analyses

To quantify differences in LMM regression slopes, we ran a two-way repeated measures ANOVA with factors visual area and stimulus conditions across participants. To quantify differences in pRF model cv-R^2^, we ran a two-way repeated measures ANOVA with factors pRF model and visual area (ROI) across voxels of all participants and visual areas. For both ANOVA results, if there was a main effect (*p*<0.05), we used Bonferroni-corrected post-hoc multiple comparison t-tests (two-sided) to evaluate differences between pRF models, or visual area and stimulus condition. We used Pearson’s correlation (*r*) to quantify the relationship between participant slopes averaged across conditions and effective pRF size, exponents (*CST_n_* or DN-ST *n*), time constants (CST_opt_ tau, DN-ST tau_1_ and tau_2_), or DN-ST semi-saturation constants across visual areas.

## Acknowledgements

We thank Brian Wandell for fruitful discussions. This research was supported by the US NIH NEI R01 EY023915 (KGS). Funders had no role in study design, data collection and analysis, or decision to submit the manuscript.

## Author contributions

E.R.K and K.G.S. designed the experiment. K.G.S. provided funding. E.R.K. and I.K. collected data and wrote computational framework. E.R.K. analyzed the data. K.G.S. oversaw computational framework and data analysis. E.R.K and K.G.S. wrote the manuscript. I.K. provided feedback on manuscript.

## 7 Supplementary Material

**Supplementary Figure 1.**
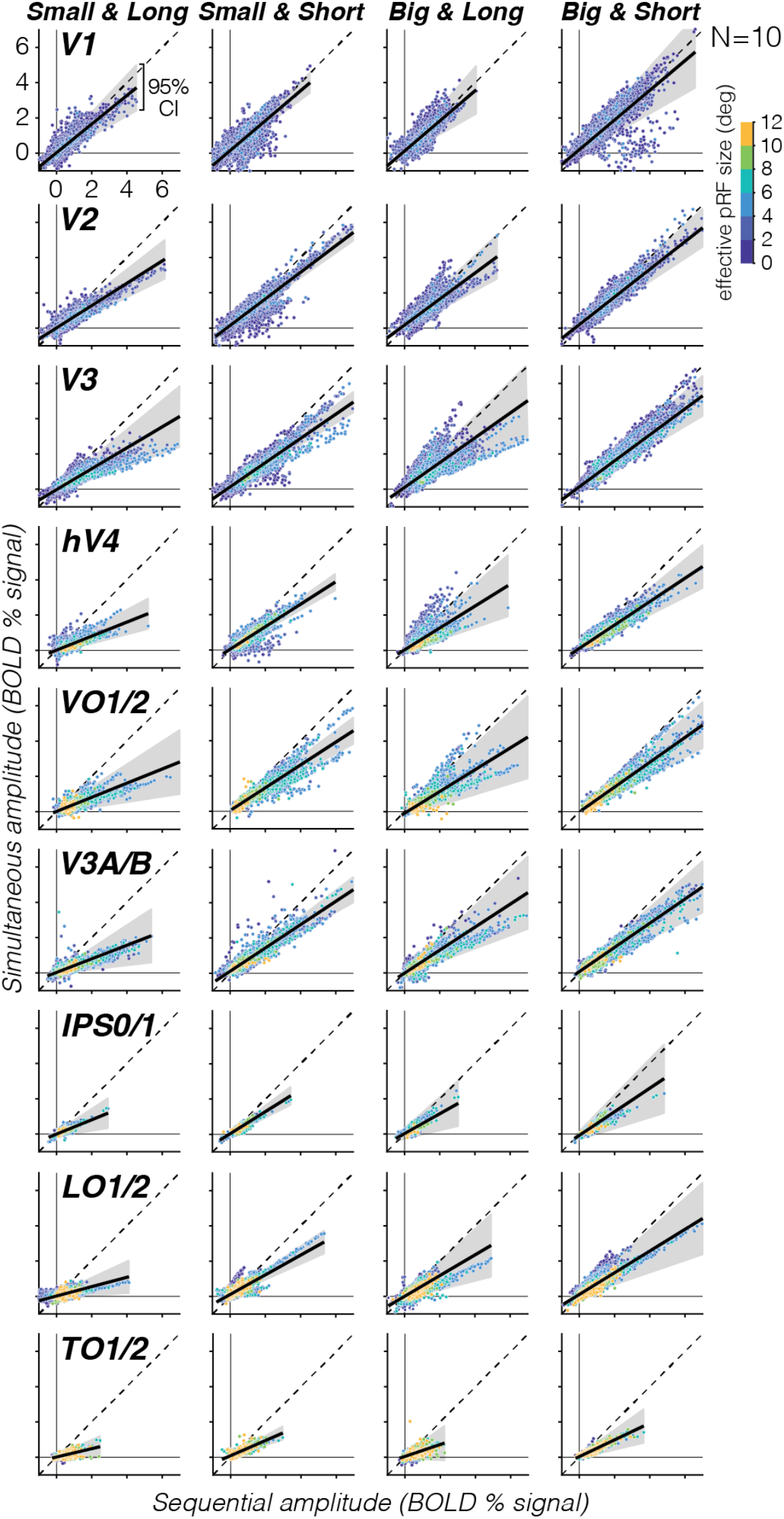
Average sequential vs simultaneous BOLD amplitude of individual voxels for all stimulus condition. Each point is a voxel are colored by effective pRF size estimated from retinotopy data in six 2° non-overlapping bins. Each panel shows data of all 10 participants. *Black solid line:* Average LMM slope across participants. *Shaded area:* 95%-confidence interval across participants. *Dashed line*: Identity line, no suppression.

**Supplementary Figure 2.**
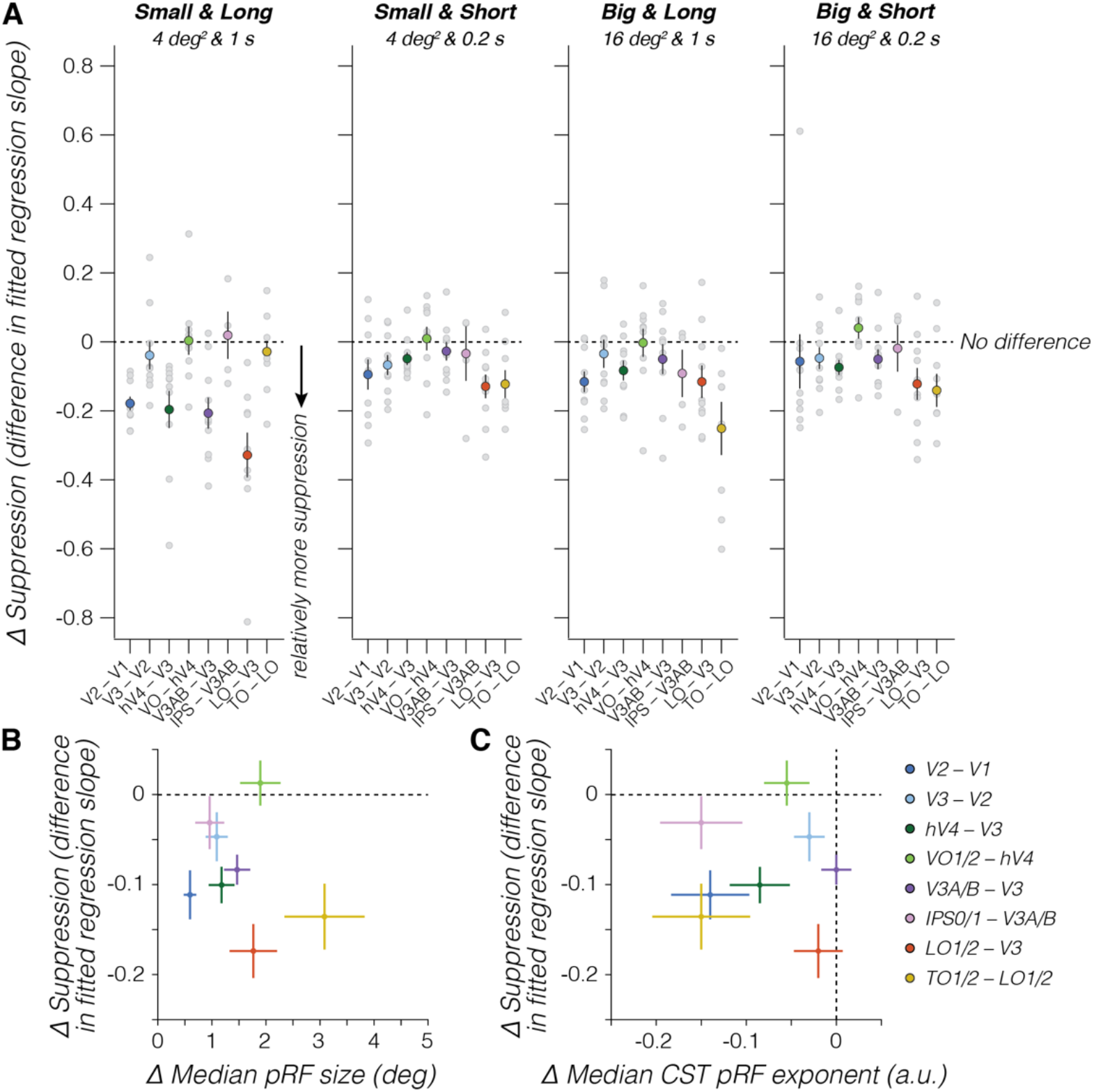
Accumulation of simultaneous suppression up the visual hierarchy. (A) Difference in suppression levels between two visual areas in the hierarchy, for each stimulus condition. Delta suppression values are derived by first subtracting participants’ regression slopes of the earlier visual area from the subsequent later visual area (e.g., V2–V1). *Large colored dots:* Group average of 10 participant’s delta suppression values for each area pair. Zero indicates no difference in suppression. Negative values indicate relatively more suppression in the later than earlier visual area. Positive values indicate relatively more suppression in the earlier than later visual area. We find that the difference in simultaneous suppression levels across consecutive visual areas varies by stimulus condition and that there is neither a monotonic increase nor plateauing of suppression across visual processing hierarchies. For example, for long duration stimuli, we observe a “sawtooth” pattern where suppression increases more from V1 to V2 than V2 to V3, and more from V3 to hV4 than hV4 to VO1/2 in the ventral stream. This suggests no monotonic increase in suppression from mid-level area hV4 to higher-level visual areas VO1/2. *Light gray dots:* Individual participant delta suppression. *Dark blue:* V2–V1. *Light blue:* V3–V2, *Dark green:* hV4–V3. *Light green:* VO1/2–-hV4. *Purple:* V3A/B–V3. *Pink:* IPS0/1 –V3A/B. *Red:* LO1/2–V3. *Yellow:* TO1/2–LO1/2. *Error bars:* SEM across 10 participants. **(B) Difference in suppression levels vs median pRF size. (C) Difference in suppression levels vs median CST pRF exponent.** In panel B and C, delta suppression values are averaged across stimulus conditions within a participant, before computing the difference in suppression between visual areas. Delta pRF size and pRF exponent are derived by first subtracting median pRF size (or CST exponent) of the earlier visual area from the later visual area within participants, then averaging differences values across 10 participants for each area pair (dots). Overall, we find no clear relation between differences in suppression levels with the difference in pRF size (B) or spatiotemporal compression (C) between paired visual areas. *Error bars*: SEM across participants. *Dashed line:* No difference in suppression slopes.

**Supplementary Figure 3.**
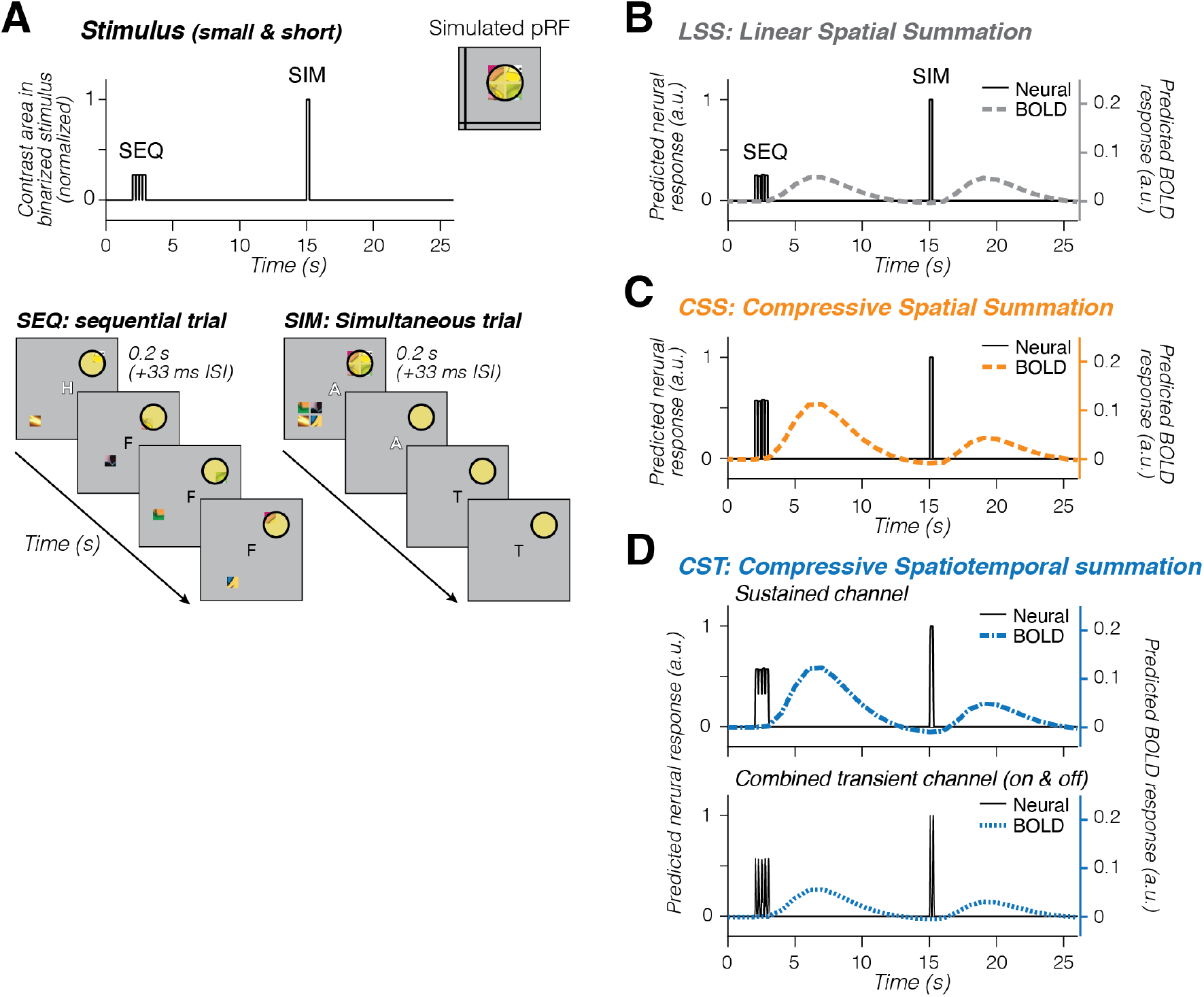
Simulated pRF model predictions for a single sequential trial followed by a simultaneous trial. Each model simulation uses a large pRF (see inset in A) that overlaps four small squares presented for 0.2 s (short square timing). **(A) Stimulus time course.** The stimulus’ visual extent is represented as the total contrast area in a binarized stimulus frame, where pixels are summed across space for each time point and normalized to set the maximum contrast area to 1. Because each trial has 4 squares per quadrant, the contrast area for each square in the sequential trial (SEQ) is a fourth of the area when all squares are shown simultaneously (SIM). **(B-D) PRF model predictions.** *Black lines & left y-axis:* predicted neural response. *Colored lines & right y-axis:* predicted BOLD response. **(B) Linear spatial summation (LSS) pRF prediction (dashed gray)**. The LSS model sums stimulus input linearly over time and space. This linearity, combined with individual squares in simultaneous and sequential trials being matched in duration and location relative to the pRF, results in the LSS model predicting no simultaneous suppression. **(C) Compressive spatial summation (CSS) pRF prediction (dashed orange).** Due to the compressive static nonlinearity, the CSS model predicts simultaneous suppression when multiple squares (simultaneously) overlap with the pRF than when a single square (sequentially) overlaps the pRF. The CSS pRF model sums linearly in time and as the overall stimulus duration is similar in our experiment between the blocks of short and long conditions, it will not predict differences in response amplitude for short vs long stimulus presentation timings (not shown in this simulation). **(D) Compressive spatiotemporal summation (CST) pRF prediction (blue)**. *Blue dot-dashed:* sustained spatiotemporal channel. *Blue dashed:* combined on- and off-transient spatiotemporal channel. By explicitly encoding neural temporal transients in milliseconds, the CST model predicts larger BOLD responses for many visual transients (SEQ) vs a few transients (SIM). The static nonlinearity produces additional subadditive spatiotemporal summation for both sustained and transient channels, including spatial subadditivity when multiple squares overlap the pRF. Consequently, both CST channels generate larger responses for sequential than simultaneous presentations, and predict different responses for the short and long conditions (not shown in simulation), which vary by a factor of 4 in number of transients (Fig 1B).

**Supplementary Figure 4.**
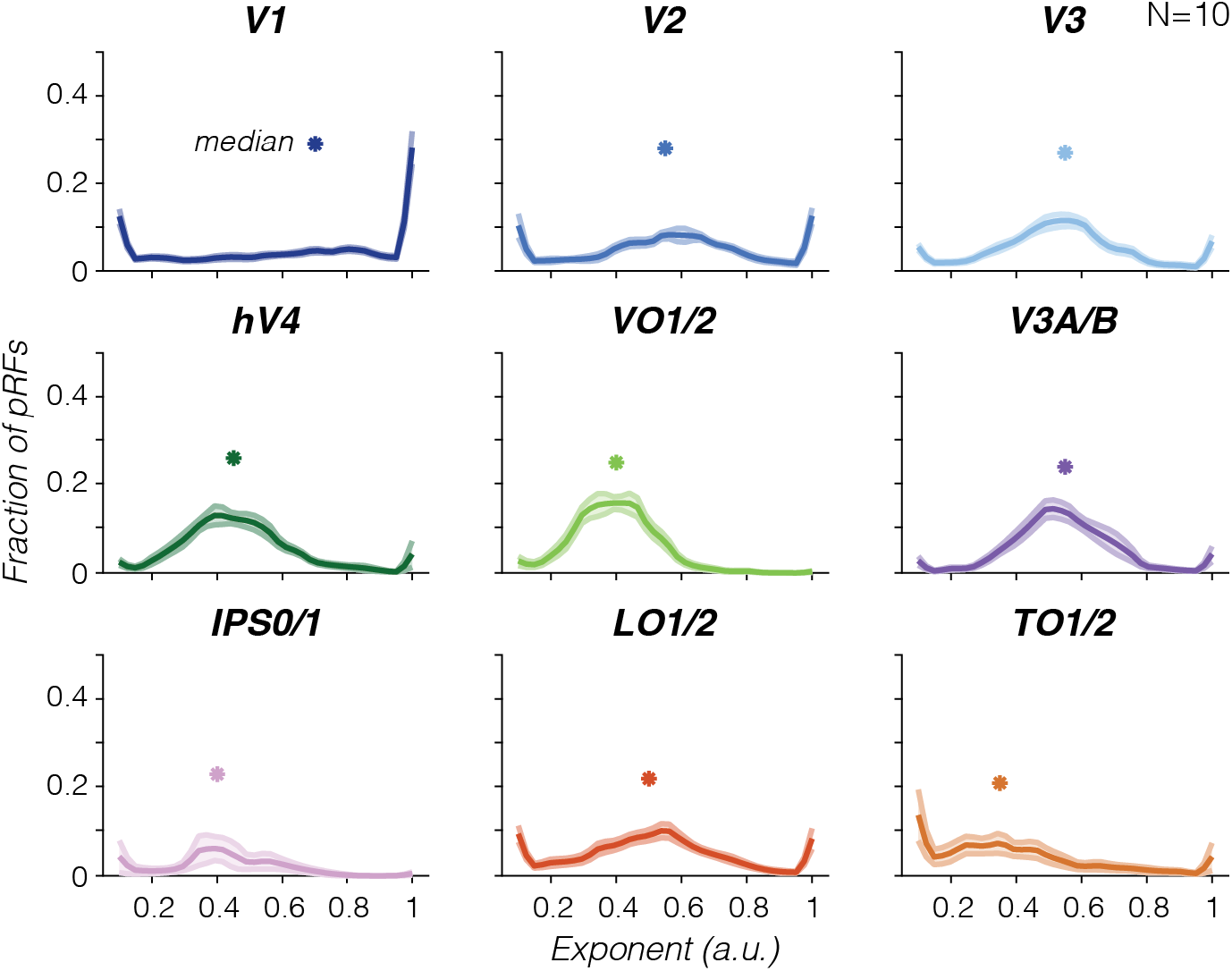
Average CST pRF exponent parameter distributions. Distributions are computed by first resampling participants’ data 1000x per visual area, then averaging distributions across participants. Both group average (line) and SEM (shaded area) of each visual area distribution are then upsampled 2x. *Asterisks:* median CST exponent value. **Eye movement analysis.** Raw horizontal and vertical gaze position (deg) and velocity (deg/s) time series of 5 participants during SEQ-SIM fMRI experiment were preprocessed as follows. First, we removed time points occurring within -100 to 100 ms of blinks. Second, given large amounts of spatial noise, we used the Identification by Two-Means Clustering algorithm^1^ to label robust fixation periods and their visual field location. If gaze locations jumped between two means due to noise, we recentered data to a single mean. Third, we removed time points (and surrounding 2 ms) if they had (i) a velocity larger than a typical saccade up to 8° (400 deg/s)^2^, (ii) an absolute gaze location that extended beyond the stimulus display (radius = 10°), or (iii) a gaze position SD 2.5x larger than SD across horizontal and vertical time series. We excluded 7 runs with < 20% data, resulting in 32 runs total. We visualized participant’s median and kernel density of gaze location across runs in visual space.

**Supplementary Figure 5.**
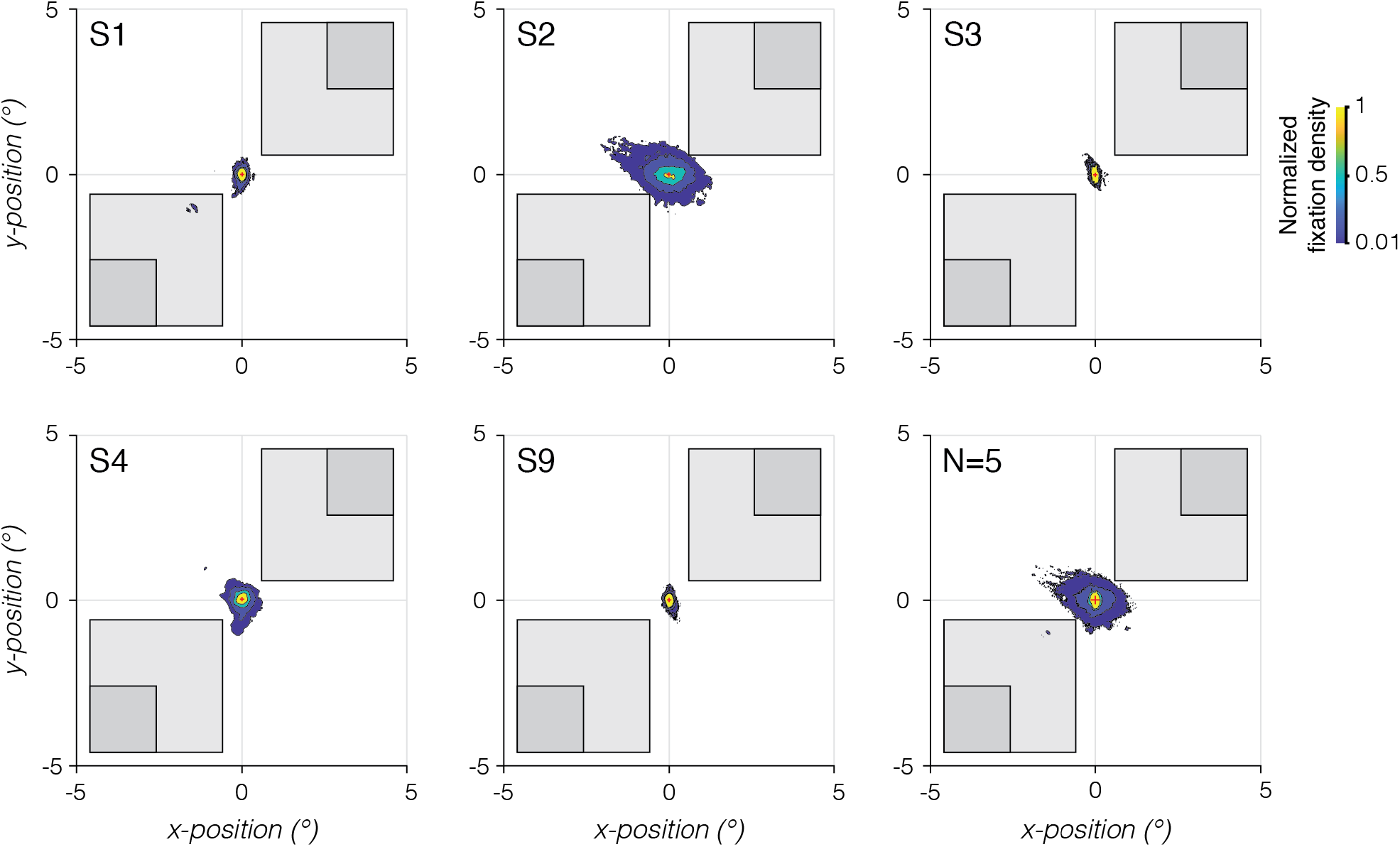
Eye fixation locations during SEQ-SIM experiment. Normalized fixation density is shown for 5 participants (S1, S2, S3, S4, S9) and across all participants (N=5). *Red cross:* Median gaze location across runs. *Contour lines*: Density at 1^st^, 10^th^, 50^th^, 100^th^ percentile, correspond to magenta, dark blue, green, and yellow sections. *Light gray squares:* Outlined location of large squares closest to fixation ([x,y]=[0,0]). *Dark gray squares:* Outlined location of small squares closest to fixation.

1. Hessels, R.S., *et al.*, Noise-robust fixation detection in eye movement data: Identification by two-means clustering (I2MC). *Behav Res Methods* **49**(5), 1802-1823 (2017).
2. Smeets, J.B. and Hooge, I.T. Nature of variability in saccades. *J Neurophysiol* **90**(1), 12-20 (2003).

**Supplementary Figure 6.**
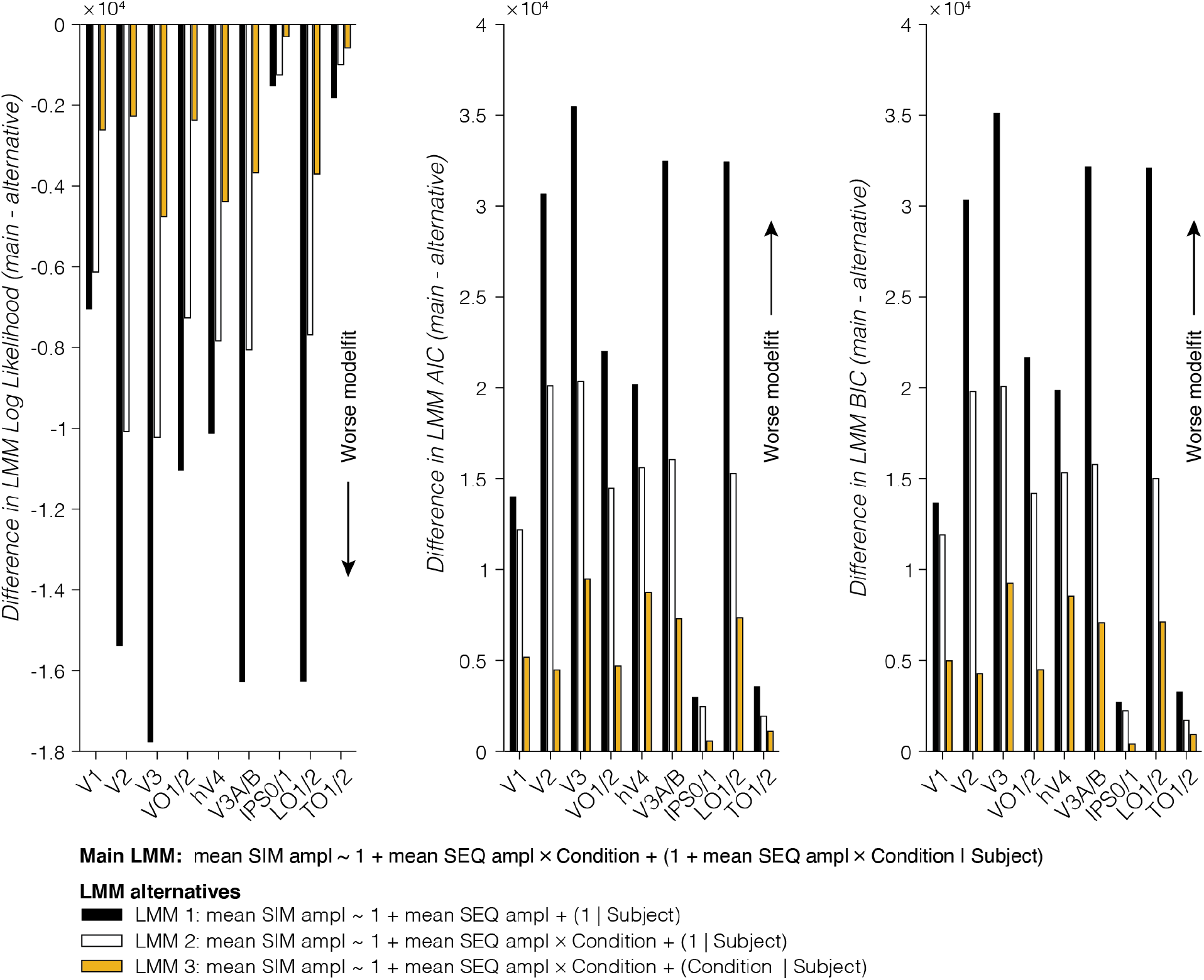
Comparison of linear mixed models (LMMs). For each model comparison metric, we computed the difference between main LMM and alternative LMM. The main LMM fits the data better than all alternative LMMs, for each visual area, on each metric. *Right:* Difference in LMM log likelihood. *Middle:* Difference in LMM AIC. *Left:* Difference in LMM BIC. The main LMM uses a fixed intercept and slope for mean sequential (SEQ) amplitude as a function of stimulus condition and allowing for random participant intercept and slope per stimulus condition. *Black bar:* Alternative LMM 1, using a fixed intercept and slope for mean sequential (SEQ) amplitude and allowing one random slope per participant. *White bar:* Alternative LMM 2, using a fixed intercept and slope for mean sequential (SEQ) amplitude as a function of stimulus condition and allowing one random slope per participant. *Yellow bar:* Alternative LMM 3, using a fixed intercept and slope for mean sequential (SEQ) amplitude as a function of stimulus condition and allowing a random slope per participant, per condition.

**Supplementary Table 1.** Summary of suppression slopes for 9 visual areas and 4 stimulus conditions. *Data are scale factors and have arbitrary units. Data are from 10 participants, except for IPS0/1 (4 participants) and TO1/2 (8 participants). M: mean. SE: standard error*.

| Visual area | Stimulus condition |  |  |  |  |  |  |  |
| --- | --- | --- | --- | --- | --- | --- | --- | --- |
|  | Small & Short |  | Small & Long |  | Big & Short |  | Big & Long |  |
|  | M | SE | M | SE | M | SE | M | SE |
| V1 | .85 | .057 | .81 | .023 | .84 | .070 | .85 | .036 |
| V2 | .75 | .028 | .63 | .023 | .78 | .037 | .78 | .055 |
| V3 | .67 | .040 | .59 | .054 | .74 | .039 | .70 | .076 |
| hV4 | .64 | .037 | .40 | .029 | .66 | .040 | .62 | .080 |
| VO1/2 | .65 | .051 | .40 | .034 | .70 | .042 | .62 | .070 |
| V3A/B | .66 | .051 | .39 | .043 | .67 | .035 | .65 | .059 |
| IPS0/1 | .63 | .061 | .41 | .054 | .67 | .10 | .56 | .056 |
| LO1/2 | .56 | .057 | 0.27 | 0.041 | .61 | .051 | .59 | .064 |
| TO1/2 | .43 | .057 | 0.24 | 0.043 | 0.47 | 0.019 | .47 | .11 |

**Supplementary Table 2.**
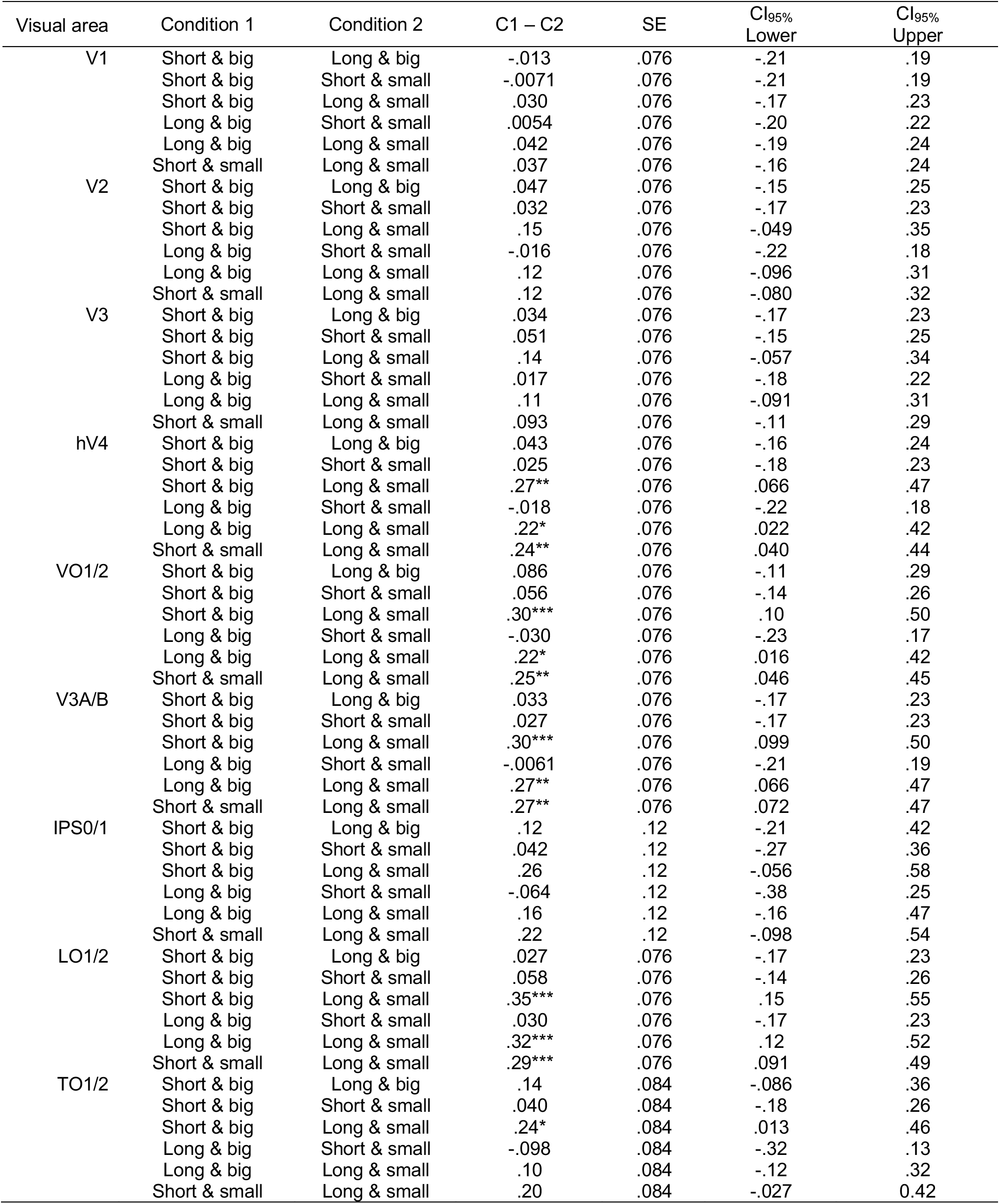
Post-hoc comparisons of suppression slopes, for each visual area and stimulus condition. Mean condition difference (C1 – C2), standard error (SE) and 95%-confidence intervals are have arbitray units (slope). P-values are Bonferroni-corrected for multiple comparisons. * *p*<0.05, ** *p*<0.01, *** *p*<0.001.

**Supplementary Table 3.**
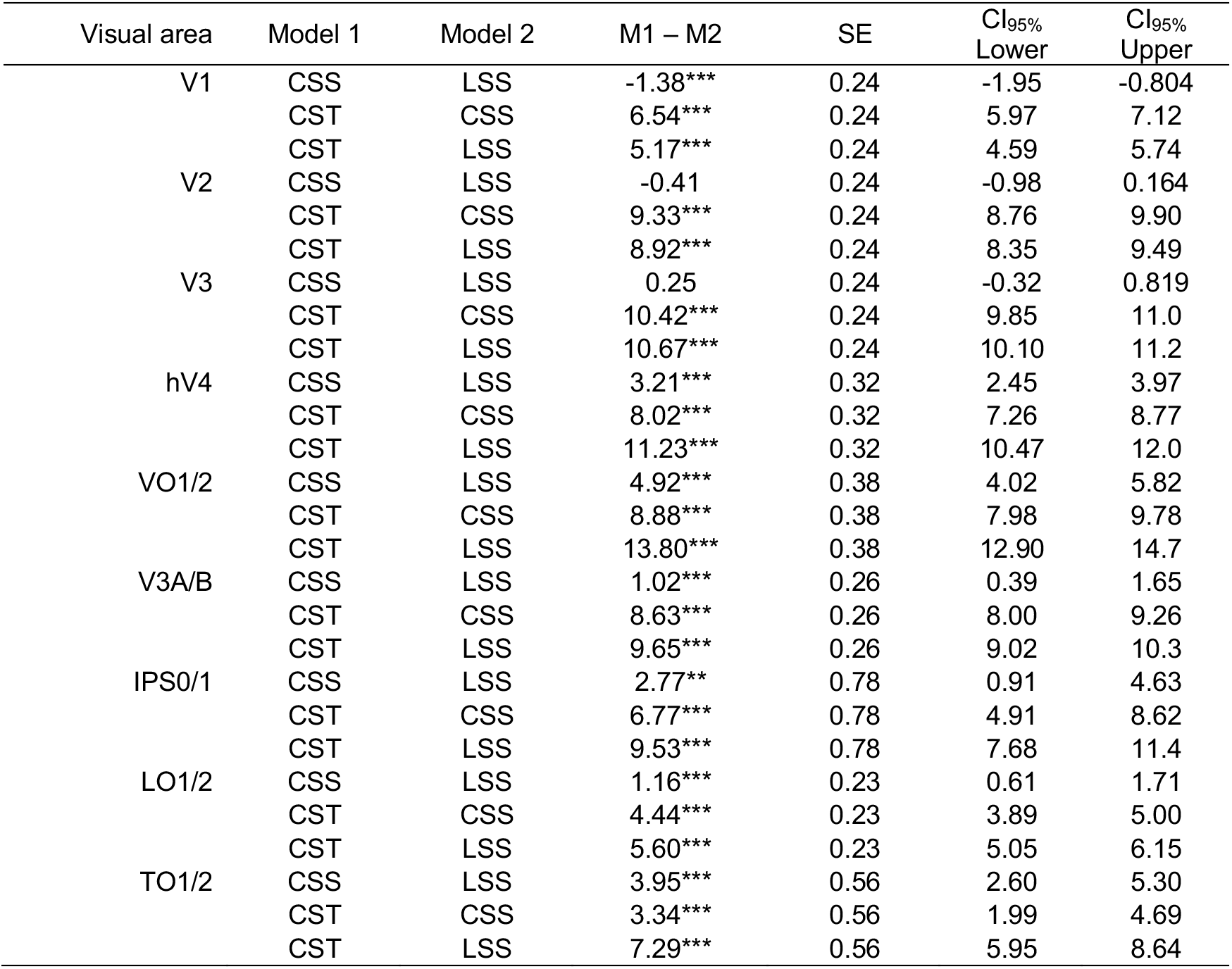
Post-hoc comparisons of pRF model performance. Mean model difference (M1 – M2), standard error and 95%-confidence intervals are in units of percent cross-validated variance explained (cv-R^2^) and correspond to violin plots in Fig 6C. P-values are Bonferroni-corrected for multiple comparisons. Significant differences are indicated with * *p*<0.05, ** *p*<0.01, or *** *p*<0.001.

**Supplementary Figure 7.**
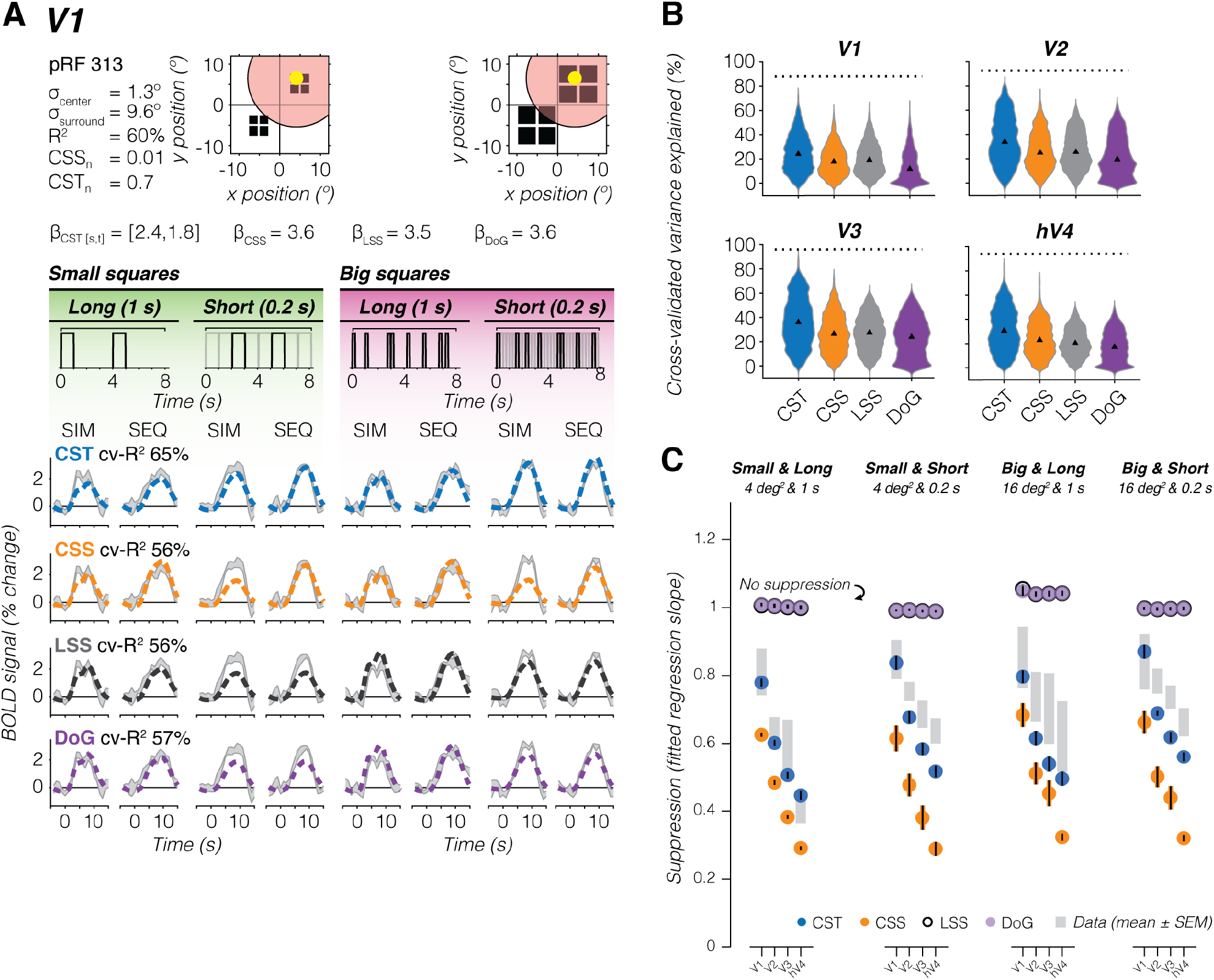
Simulated Difference of Gaussians pRF model performs similarly to the LSS pRF model. **(A) V1 example voxel.** *Gray shaded area:* Average ± SEM voxel time series, repeated for each row. PRF model fits are shown in dashed lines. *Dark Blue:* Compressive spatiotemporal summation model with fixed voxel parameters (CST, first row). *Orange:* Compressive spatial summation model (CSS, second row). *Black:* Linear spatal summation model (LSS, third row). *Purple:* Difference of Gaussians model (DoG, fourth row). DoG pRF surround size is based on each visual area’s average pRF center/surround ratio reported by Aqil *et al.* (2021)^50^. **(B) Distribution of voxel-level cross-validated variance explained for each pRF model.** *Triangle:* median. *Dotted line:* noise ceiling computed from voxel’s maximum split-half reliability across participants. CST (blue), CSS (orange), and LSS (gray) violin plots are the same as in Fig 6C. *Purple*: DoG. Each participant’s data is resampled 1000x for each visual area. **(C) Model-based prediction of simultaneous suppression vs observed simultaneous suppresion.** Panel shows the same observed and predicted CST, CSS, and LSS suppression levels as Fig 7, with the addition of predicted suppression levels by the simulated DoG pRFs (semi-saturated filled purple circles) Model-based points and errorbars show average and SEM across 10 participants.

**Supplementary Figure 8.**
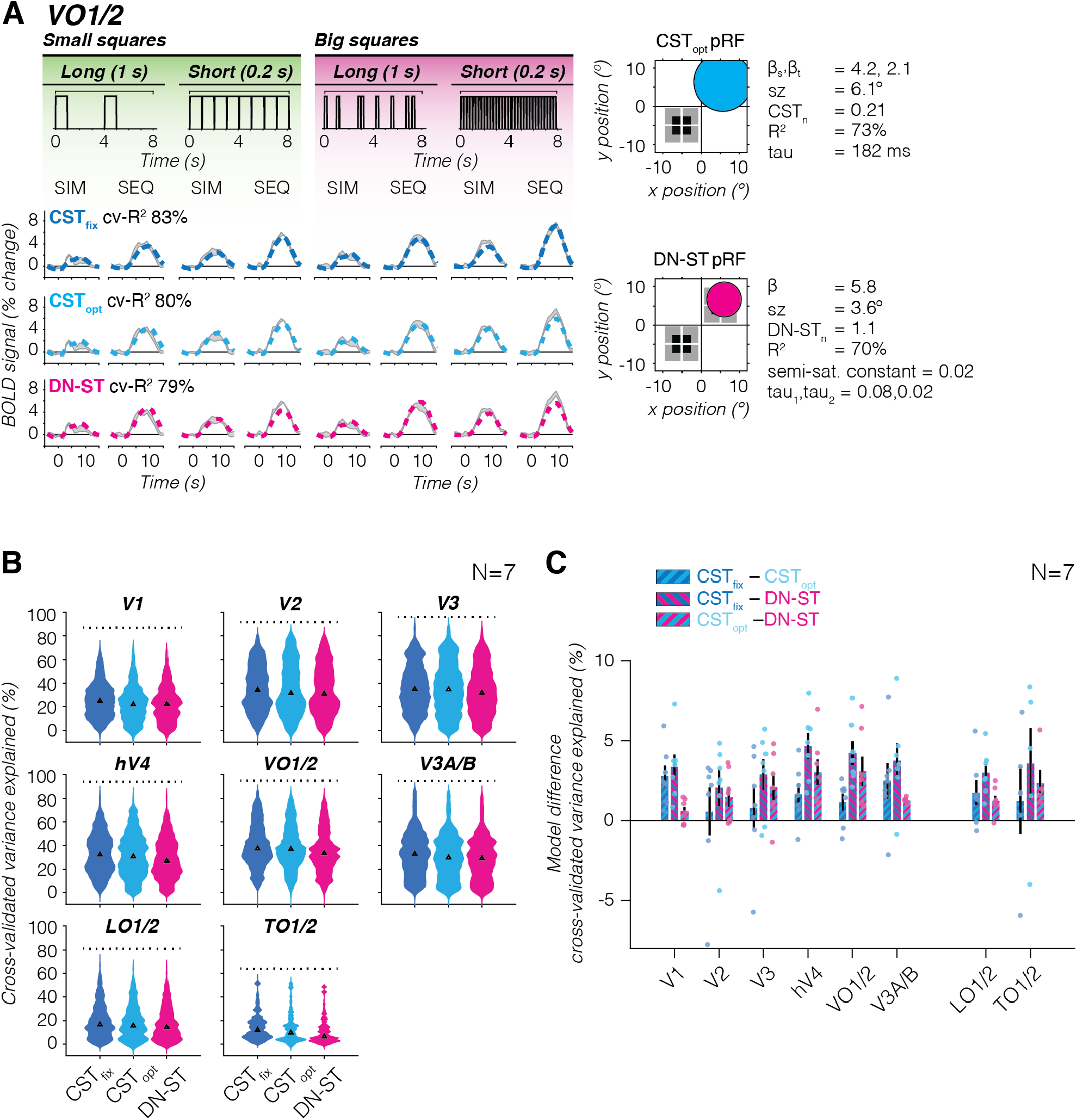
Comparison of fixed vs optimized CST and DN-ST pRF model performance. **(A) VO1/2 example voxel.** *Gray shaded area:* Average ± SEM voxel time series. CST_fix_ data are from the same voxel as in Fig 6B. CST_opt_ and DN-ST example voxels show corresponding time series to CST_fix_ time series from the same participant (S3), but have different pRF parameters from Kim *et al.* (2024)^51^. PRF model fits are shown in dashed lines. *Dark Blue:* Compressive spatiotemporal summation model with fixed voxel parameters (CST_fix_, top row). *Cyan:* Compressive spatiotemporal summation model with optimized voxel parameters (CST_opt_, middle row). *Pink:* Divisive normalization spatiotemporal summation model with optimized voxel parameters (DN-ST, bottom row). **(B) Distribution of voxel-level cross-validated variance explained for each pRF model.** Each violin plot contains data from the 7 overlapping participants with Kim *et al.* (2024). *Triangle:* Median. *Dotted line:* Noise ceiling computed from voxel’s maximum split-half reliability across participants. *Blue:* CST_fix_. *Cyan*: CST_opt_. *Pink:* DN-ST. Each participant’s data is resampled 1000x for each visual area. A two-way repeated measures ANOVA revealed significant effects of pRF model (*F*(2)=1.1×10^2^, *p*=6.3×10^-47^) and ROI (*F*(7)=1.3×10^3^, *p*<10^-47^) on cv-R^2^, with CST models outperforming DN-ST on average by about 3%, as well as a significant interaction between pRF model and ROI (*F*(2,7)=5.0, *p*=1.6×10^-9^). Post-hoc Bonferroni-corrected t-tests revealed that both CST models significantly explain more cv-R^2^ than the DN-ST model in all visual areas, except for CST_opt_ in TO1/2. In addition, the CST_fix_ model explains significantly more cv-R^2^ than the CST_opt_ in several visual areas (V1, hV4, V3A/B, LO1/2, and TO1/2). **(C) Pairwise model comparison for each visual area.** *Bars:* Group average voxelwise difference in cv-R^2^ between two pRF models. *Error bars:* SEM across participants. *Individual dots:* Average difference for each participant. *Dark blue–cyan:* CST_fix_ vs CST_opt_. *Dark blue–pink:* CST_fix_ vs DN-ST. *Cyan–pink:* CST_opt_ vs DN-ST.

**Supplementary Table 4.** Post-hoc comparisons of spatiotemporal pRF model performance. Mean model difference (M1 – M2), standard error, and 95%-confidence intervals are in units of percent cross-validated variance explained (cv-R^2^) and correspond to violin plots in Supplementary Fig 8B. Data are from 7 overlapping participants with Kim *et al.* (2024)^51^. IPS0/1 results are removed as only two participants contributed data for these visual areas. P-values are Bonferroni-corrected for multiple comparisons. Significant differences are indicated with * *p*<0.05, ** *p*<0.01, or *** *p*<0.001.

| Visual area | Model 1 | Model 2 | M1 – M2 | SE | CI<sub>95%</sub><br>Lower | CI<sub>95%</sub><br>Upper |
| --- | --- | --- | --- | --- | --- | --- |
| V1 | CST<sub>fix</sub> | CST<sub>opt</sub> | 2.35*** | 0.55 | 1.03 | 3.67 |
|  | CST<sub>fix</sub> | DN-ST | 2.89*** | 0.55 | 1.58 | 4.21 |
|  | CST<sub>opt</sub> | DN-ST | 0.54 | 0.55 | -0.77 | 1.86 |
| V2 | CST<sub>fix</sub> | CST<sub>opt</sub> | -0.16 | 0.43 | -1.18 | 0.870 |
|  | CST<sub>fix</sub> | DN-ST | 1.24* | 0.43 | 0.21 | 2.26 |
|  | CST<sub>opt</sub> | DN-ST | 1.39** | 0.43 | 0.36 | 2.42 |
| V3 | CST<sub>fix</sub> | CST<sub>opt</sub> | 0.95 | 0.44 | -0.10 | 1.99 |
|  | CST<sub>fix</sub> | DN-ST | 3.00*** | 0.44 | 1.96 | 4.05 |
|  | CST<sub>opt</sub> | DN-ST | 2.06*** | 0.44 | 1.01 | 3.10 |
| hV4 | CST<sub>fix</sub> | CST<sub>opt</sub> | 1.79* | 0.64 | 0.27 | 3.31 |
|  | CST<sub>fix</sub> | DN-ST | 5.08*** | 0.64 | 3.56 | 6.60 |
|  | CST<sub>opt</sub> | DN-ST | 3.23*** | 0.64 | 1.76 | 4.81 |
| VO1/2 | CST<sub>fix</sub> | CST<sub>opt</sub> | 0.92 | 0.63 | -0.58 | 2.43 |
|  | CST<sub>fix</sub> | DN-ST | 4.56*** | 0.63 | 3.05 | 6.07 |
|  | CST<sub>opt</sub> | DN-ST | 3.63*** | 0.63 | 2.12 | 5.15 |
| V3A/B | CST<sub>fix</sub> | CST<sub>opt</sub> | 3.34*** | 0.51 | 2.12 | 4.56 |
|  | CST<sub>fix</sub> | DN-ST | 4.58*** | 0.51 | 3.36 | 5.80 |
|  | CST<sub>opt</sub> | DN-ST | 1.24* | 0.51 | 0.017 | 2.46 |
| LO1/2 | CST<sub>fix</sub> | CST<sub>opt</sub> | 1.35* | 0.53 | 0.088 | 2.61 |
|  | CST<sub>fix</sub> | DN-ST | 2.88*** | 0.53 | 1.62 | 4.14 |
|  | CST<sub>opt</sub> | DN-ST | 1.53* | 0.53 | 0.27 | 2.79 |
| TO1/2 | CST<sub>fix</sub> | CST<sub>opt</sub> | 4.40* | 1.56 | 0.65 | 8.14 |
|  | CST<sub>fix</sub> | DN-ST | 6.61*** | 1.56 | 2.86 | 10.4 |
|  | CST<sub>opt</sub> | DN-ST | 2.21 | 1.57 | -1.55 | 5.98 |

**Supplementary Figure 9.**
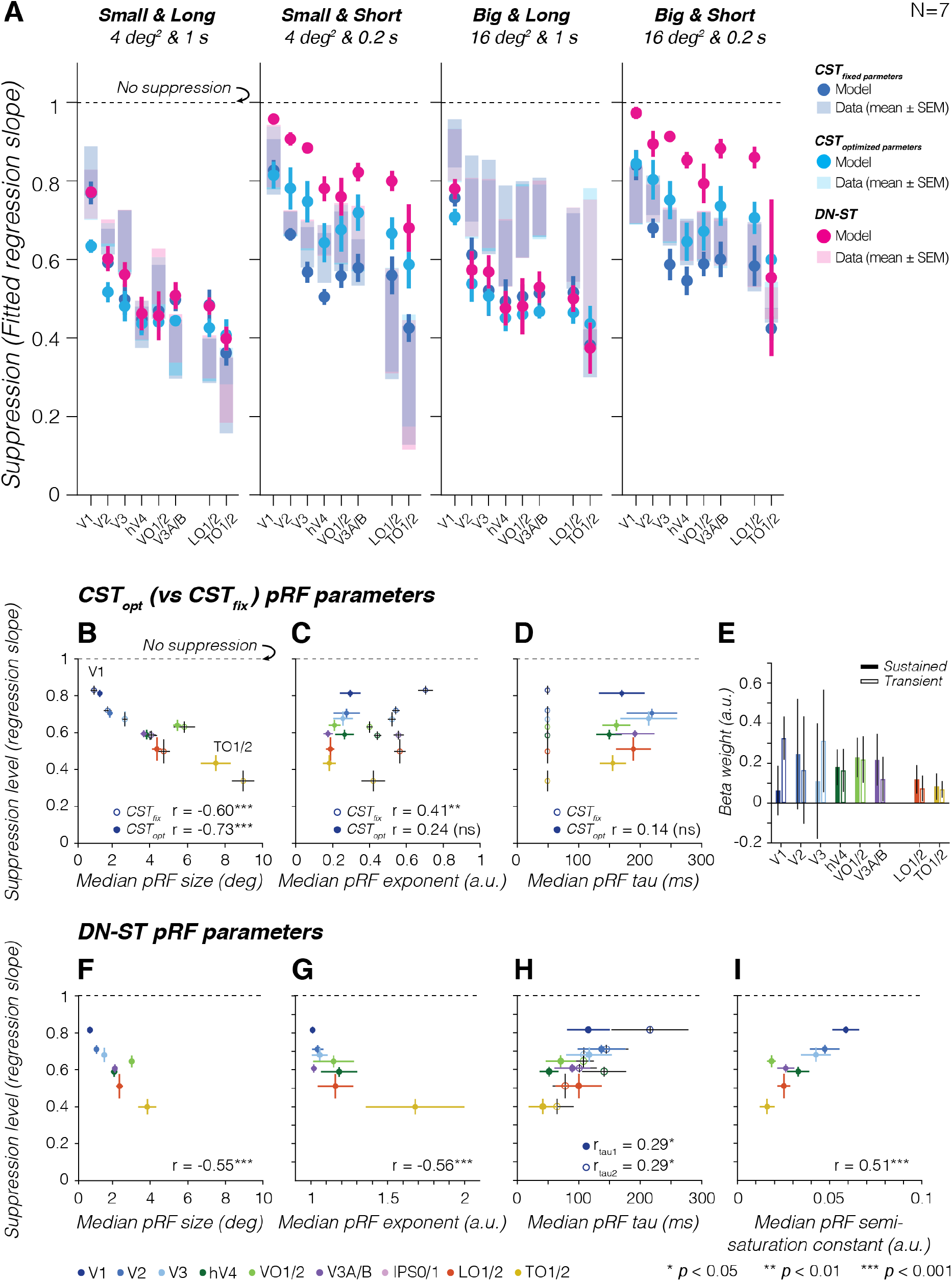
Comparison of compressive spatiotemporal pRF models. Suppression levels and pRF parameters are from voxels overlapping SEQ-SIM squares for the 7 participants overlapping with Kim *et al.* (2024)^51^. In all panels: *dots/bars* show average across 7 participants. *Error bars:* SEM across 7 participants. IPS0/1 results are removed as only two participants contributed data for these visual areas. **(A) CST_fix_, CST_opt_, and DN-ST pRF model-based predictions of simultaneous suppression**. Shaded *dark blue bars:* Average ± SEM observed suppression levels in voxels with CST_fix_ pRFs. *Shaded cyan bars:* Average ± SEM observed suppression levels in voxels with CST_opt_ pRFs. *Shaded pink bars:* Average ± SEM observed suppression levels in voxels with DN-ST pRFs. *Dark blue circles*: Average ± SEM predicted simultaneous suppression by CST_fix_ pRFs. *Cyan filled circles:* Average ± SEM predicted simultaneous suppression by CST_opt_ pRFs. *Pink filled circles:* Average ± SEM predicted simultaneous suppression by DN-ST pRFs. The DN-ST pRFs tends to underpredict simultaneous suppression levels for short stimulus durations, which may be due to less accurate temporal parameter recovery for the DN-ST compared to the CST model^51^ and/or to the DN-ST model being more sensitive to prolonged stimulus durations than visual transients^40,42^ whereas the SEQ-SIM experiment is dominated by latter. **(B-D). Simultaneous suppression levels vs CST_opt_ and CST_fix_ pRF parameters.** For panels B-D (effective size, exponent, and tau pRF parameters), we first computed the median across pRFs of a visual area for each participant (similar to Fig 8), then we calculated the average of this median value across 7 participants. Pearson’s correlation (*r*) is computed using individual participant data. *Filled colored circles*: CST_opt_ pRF parameters. *White circles with colored outline:* CST_fix_ pRF parameters. (B) Simultaneous suppression level vs median CST_opt_ pRF effective size. (C) Simultaneous suppression level vs CST_opt_ exponent (static nonlinearity). (D) Simultaneous suppression level vs time constant τ (tau). We find that the range of estimated spatiotemporal compression (exponent) in the CST_opt_ pRF model in the Kim *et al.* (2024) experiment is smaller, and compression is overall stronger than the spatiotemporal compression in the CST_fix_ pRF model, which was optimized to the SEQ-SIM data. **(E) Average β-weights of sustained and transient channels for CST_opt_ pRF model.** Beta weights are averaged first within a participant’s visual area, then averaged across participants per visual area. *Colored bars:* Sustained channel. *White bars:* Combined transient channel. Channels do not differ significantly from one another across visual areas. **(F-I) Simultaneous suppression levels vs DN-ST pRF parameters.** For all panels, we first computed the median across pRFs of a visual area for each participant (similar to Fig 8), then we calculated the average median value across participants. Pearson’s correlation (*r*) is computed using individual participant data. (F) Simultaneous suppression level vs median DN-ST pRF effective size. (G) Simultaneous suppression level vs exponent (DN-ST *n*). (H) Simultaneous suppression level vs time constants. *Filled colored circles*: τ_1_ parameter, time constants of the IRF. A smaller tau_1_ results in an IRF that peaks earlier. *White circles with colored outline:* τ_2_ parameter, time constant of the exponential decay. A smaller τ_2_ results in quicker decay and stronger compression. (I) Simultaneous suppression level vs semi-saturation constant (*σ_DN_*). A smaller semi-saturation results in stronger compression within the pRF. Significant effects are indicated with *\* p*<0.05, \**\* p*<0.01 or *** *p*<0.001; *ns*: not significant.

## Notes

### Competing Interest Statement

The authors have declared no competing interest.

### Summary of Updates

We clarified our reasoning in the introduction, results, and discussion. We expanded the methods section, and added multiple new analyses: including three additional population receptive field modelfits, and new supplementary figures 2, 7-9.

